# Proteostasis sustains T cell differentiation potential and tumor-infiltrating lymphocyte function

**DOI:** 10.64898/2026.02.08.704716

**Authors:** Nicole E Scharping, Xuezhen Ge, Maria Inês Matias, Fulin Jiang, Allison Cafferata, Maximilian Heeg, Alexander Monell, Giovanni Galletti, Kitty P Cheung, Angelica Rock, Nick Thao, Sydnye L Shuttleworth, Michael A. Bauer, Kennidy K Takehara, Amir Ferry, Sara Quon, Brian Koss, Samuel A Myers, Eric J Bennett, Ananda W Goldrath

## Abstract

Tumor-infiltrating lymphocytes (TIL) often fail to restrain tumor growth due to progressive differentiation to an ‘exhausted’ state. In healthy tissues, tissue-resident memory T cells (T_RM_) maintain protection for years, and patient tumors that contain TIL with T_RM_ features are associated with better prognosis. Proteomic and transcriptomic profiling of T cell populations identified proteostasis as a significant factor distinguishing T_RM_ and progenitor-exhausted TIL from terminally-exhausted TIL, including loss of E3 ubiquitin ligases NEURL3, RNF149, and WSB1, with accumulation of unfolded proteins in spite of functional proteasome activity. Enforced expression of these ligases by TIL preserved stem-like TCF1^+^ populations and improved anti-tumor function, whereas their knockout impaired TIL and altered T cell differentiation in acute infection. Sustained ligase expression rescued accumulation of unfolded proteins in TIL and improved immunotherapy outcome in preclinical models, highlighting the critical role of proteostasis in TIL function and identifying new avenues for advancing cancer immunotherapy.

## Introduction

Naive CD8^+^ T cell activation involves the integration of diverse signals to shape cell fates^1^. In response to tumor or chronic viral infection, persistent TCR stimulation drives T cell differentiation through a progressive loss of function^2^, with a spectrum of functionality from more stem-like T_PEX_ to terminally-differentiated T_EX_^3,4^. T_PEX_ retain self-renewal capacity via TCF1 expression and give rise to T_EX_. T_EX_ are characterized by co-inhibitory receptor expression including PD-1 and TIM-3, metabolic dysfunction with high levels of reactive oxygen species production from dysfunctional mitochondria, an altered chromatin landscape associated with expression of the transcription factor TOX that contributes the T_EX_ epigenomic state, and limited cytokine polyfunctionality^5–7^.

The T_RM_ population is comprised of long-lived memory T cells that are essential for surveying barrier and non-lymphoid tissues, by providing a first-line defense against reinfection^8^. A population of TIL with T_RM_ features that are distinct from T_EX_ and T_PEX_ has also been identified in some patient tumors^4,9^. We have previously shown that enforcing adaptations consistent with tissue residency through the modulation of transcription factors or immunometabolic pathways are sufficient to improve tumor control in mice^10–12^. In humans, cancer patients with T_RM_-like TIL show increased overall survival and have improved outcomes after immunotherapeutic checkpoint blockade^9,13^. Thus, elucidating the relationships among T_PEX_, T_EX_, and T_RM_-like TIL populations is important to improve our knowledge of T cell biology and to facilitate the development of cancer immunotherapies.

Protein homeostasis (proteostasis) is an underexplored facet of T cell exhaustion, despite its known function in maintaining stem-like cell states. Targeted protein degradation plays a crucial role in regulating stem cell differentiation and embryonic development via the ubiquitin-proteasome system (UPS), regulating protein turnover by appending chains of ubiquitin onto target proteins^14–16^. E3 ubiquitin ligases are the substrate-specifying arm of the UPS which catalyzes ubiquitin transfer onto target proteins, leading to proteasome-dependent degradation. Beyond its role in cellular differentiation, proteostasis is critical for limiting excess protein accumulation, thus decreasing cellular stress. Accumulation of unfolded proteins due to rapid proliferation or metabolic dysfunction can trigger endoplasmic reticulum (ER) stress and the unfolded protein response (UPR)^17,18^ Furthermore, E3-dependent protein degradation is required for optimal responses to environmental stresses, such as hypoxia and increased reactive oxygen species^19^ and chronic stress can result in the accumulation of unfolded proteins that drive cellular dysfunction^20^.

For T cells, some aspects of proteostasis have been characterized; in naive T cells, activation increases protein synthesis to meet the energy demands of rapid proliferation and differentiation^21–23^. Regulation of protein turnover has been characterized in effector T cell differentiation, as proteasome function plays a role in directing cell fates and inhibition during T cell proliferation has been shown to halt cell-cycle progression^24–26^. Memory T cells maintain a high level of protein turnover at steady state and rely on proteasomal activity for development and survival^27–29^. While more than 500 genes encoding E3 ligases have been identified in the human genome^16^, the roles of specific E3 ligases in immune cells are understudied. A well-characterized example in many cell types, including T cells, is the E3 ubiquitin ligase Von Hippel-Lindau (VHL), which targets the transcription factor hypoxia-inducible factor (including HIF1α and HIF2α), allowing the cells to respond transcriptionally to local oxygen levels^30^. We have previously shown that the loss of VHL in T cells increases HIF activity, promoting T cell activity in infections and tumors^12,31^. ^32–36^However, which of the hundreds of T cell-expressed ligases are required to maintain proteostasis through regulated protein degradation, and how ligase dysfunction impacts T cell function, are largely unknown.

Here, we show that modulation of T_RM_- and T_PEX_-associated E3 ubiquitin ligases NEURL3, RNF149, and WSB1 alters T cell function. Ligase overexpression in TIL increases TCF1^+^ T cell accumulation and improves T cell function, while ligase knockout (KO) in T cells responding to cancer or acute infection alters T cell differentiation. We utilize low-input mass spectrometry for unbiased proteomics, enabling for the identification of thousands of proteins from *ex vivo*-sorted T cells, including T_PEX_ and T_EX_. In characterizing T_EX_, we show that despite possessing functional proteasomes, T_EX_ proteostasis is negatively impacted by unfolded protein accumulation and by targeting upstream E3 ligase activity, it is possible to rescue this unfolded protein accumulation. In addition, we enhance proteostasis via ligase overexpression in combination with checkpoint blockade to boost the efficacy of adoptive cell therapy. Together, these data demonstrate the role of proteostasis in T cell exhaustion and identify a new avenue for immunotherapy.

## Results

### Rescue of TIL function by sustained expression of T_RM_-associated E3 ubiquitin ligases NEURL3, RNF149 and WSB1

To understand the transcriptional relationships between T_RM_ and TIL populations, we first analyzed enrichment of tissue-residency-associated genes among endogenous T_PEX_ cells (CD8^+^PD1^lo/int^Tim3^−^TCF1^hi^TOX^lo^) and T_EX_ cells (CD8^+^PD1^hi^Tim3^+^TCF1^lo^TOX^hi^) sorted from day 14 murine B16 melanoma tumors (Figure 1A, Supplement Figure 2A, Supplemental Table 1)^32,33^. This comparison revealed almost two-thirds of the T_RM_-associated genes were expressed in T_PEX_, but downregulated or repressed in T_EX_. Of the T_RM_-associated genes lost with terminal-differentiation to T_EX_, 24% were associated with protein folding and degradation, including genes encoding for E3 ubiquitin ligases neuralized E3 ubiquitin protein ligase 3 (*Neurl3*, also known as *Lincr*), RING finger protein 149 (*Rnf149*), and WD repeat and suppressor of cytokine signaling (SOCS) box-containing 1 (*Wsb1*) (Figure 1B). NEURL3 contains a neuralized domain, and based on sequence-structure annotation, both NEURL3 and RNF149 contain RING domains, which facilitate ubiquitin transfer from E2 ubiquitin-conjugating proteins to target proteins^34,35^. WSB1 contains a SOCS domain (similar to VHL) and can act as a substrate adaptor protein for the multisubunit CUL5 RING ligase (CRL5) complex to facilitate ubiquitin transfer to substrate proteins^36,37^. However, the functions of these three ligases in immune cells are unknown.

**Figure 1:**
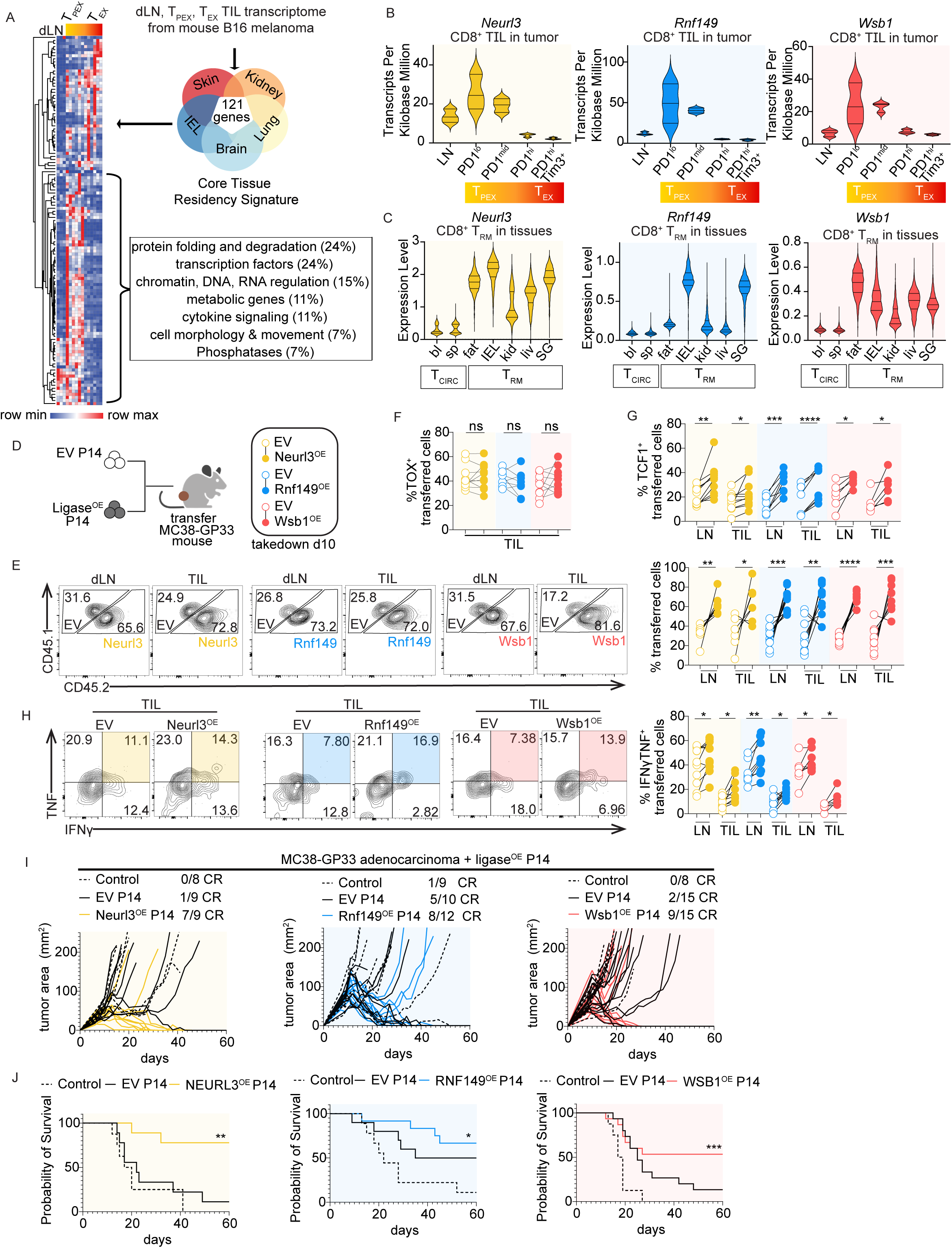
E3 ubiquitin ligases NEURL3, RNF149 and WSB1 are expressed in T_PEX_ and T_RM_, and sustained expression can rescue TIL function in MC38 adenocarcinoma. A) Heatmap of bulk RNA-seq from endogenous CD8^+^ T cells sorted from B16 mouse melanoma dLN and TIL (GSE175408^33^). TIL sorted on PD1^lo^, PD1^mid^, PD1^hi^, PD1^hi^Tim3^+^, data were filtered using the core T_RM_ gene signature^32^ and quantified via hierarchical clustering by rows and columns using one minus Pearson correlation (Supplemental Table 1). B) *Neurl3*, *Rnf149*, or *Wsb1* RNA TPM (GSE175408^33^). C) *Neurl3*, *Rnf149*, and *Wsb1* RNA expression levels from scRNA-seq of CD8^+^ T_RM_ cells isolated from the IEL, kidney, SG, fat and liver, and memory from the spleen and blood (GSE182275^78^), quantified after MAGIC imputation. D) Experimental schematic: Congenically mismatched P14 T cells were activated, transduced at 24 h with Empty Vector (EV) control or ligase-overexpressing retrovirus (Neurl3^OE^, Rnf149^OE^, or Wsb1^OE^). Next day, cells were sorted on ametrine reporter expression, then cultured for three additional days, mixed 50:50 (250,000 EV + Ligase^OE^; 500,000 total), then transferred into d10 MC38-GP_33–41_-bearing mice; dLN and TIL were analyzed 10 days later (BioRender). E) Representative flow plots and quantification of transferred P14 T cell frequencies as in D (*Neurl3*^OE^ n=7, *Rnf149*^OE^ n=11, *Wsb1*^OE^ n=10); three independent experiments. F) Tox^+^ P14 T cell frequencies in tumors as in D (*Neurl3*^OE^ n=10, *Rnf149*^OE^ n=8, *Wsb1*^OE^ n=9); three independent experiments. G) TCF1^+^ P14 T cell frequencies as in D (*Neurl3*^OE^ n=10, *Rnf149*^OE^ n=8, *Wsb1*^OE^ n=6); two or more independent experiments. H) Representative flow plots and quantification of IFNɣ^+^TNF^+^ P14 T cell frequencies as in D followed by 4 h of peptide restimulation plus protein transport inhibitor (*Neurl3*^OE^ n=10, *Rnf149*^OE^ n=8, *Wsb1*^OE^ n=7); three independent experiments. I) Quantification of MC38-GP_33–41_ tumor area in mice that received no T cells, EV P14 or *Neurl3*^OE^, *Rnf149*^OE^, or *Wsb1*^OE^ P14 T cells on d8 after tumor cell implantation. CR = complete regression; two independent experiments. J) Survival data from I. Statistical significance was calculated using paired Student’s *t*-tests (E-H), or log-rank (Mantel-Cox) test (J). Connecting lines indicate that cells were isolated from the same mouse.

We found that antigen-specific T_RM_ isolated from multiple tissues expressed *Neurl3*, *Rnf149*, and *Wsb1* (Figure 1C). In a single-cell (sc) RNA-seq time-course of P14 T cells responding to the lymphocytic choriomeningitis virus (LCMV) Armstrong, ligase expression in the spleen and small intestine intraepithelial lymphocytes (IEL) was observed on day 5 (d5) of infection and peaked at effector to early memory timepoints (Supplemental Figure 2B). *In vitro* exhaustion also led to significantly decreased ligase expression, an observation consistent with data from *ex vivo* T_EX_ (Supplemental Figure 2C). To determine if enforced ligase expression could improve TIL function, we overexpressed NEURL3, RNF149, or WSB1 in P14 TCR-transgenic CD8^+^ T cells, specific for the LCMV glycoprotein 33–41 peptide (GP_33–41_) (Supplemental Figure 2D). Ligase-expressing T cells were co-adoptively transferred with an equal number of empty vector (EV) congenically-mismatched P14 T cells into MC38 adenocarcinoma-bearing mice that expressed GP_33–41_ (Figure 1D, Supplemental Figures 1A). After 10 days, ligase-overexpressing P14 T cells were more abundant in the tumor-draining lymph node (dLN) and the tumor, had a higher frequency of TCF1-expressing cells, and were more polyfunctional following *ex vivo* GP_33–41_ peptide restimulation compared to EV T cells (Figure 1E-H). However, there were no differences in the frequencies of Tim3^hi^Tcf1^lo^ cells or TOX^+^ cells between control and ligase-overexpressing TIL (Figure 1F, Supplemental Figure 2E). Ligase-overexpressing cells also showed enhanced accumulation and improved TIL function in B16-GP_33–41_ melanoma-bearing mice, but only dLN P14 displayed an increased frequency of TCF1-expressing cells (Supplemental Figure 2F-J). This phenotype indicated that ligase overexpression supported a stem-like population in the dLN, where this phenotype was more apparent in response to a highly immunogenic tumor (MC38) compared to a poorly immunogenic tumor (B16)^38^. To examine the functional impact of ligase overexpression, we used cell therapy-sensitive MC38-GP_33–41_ where EV- or ligase-transduced P14 T cells were transferred into mice after tumor implantation, and tumor growth was monitored (Figure 1I). All three ligase-overexpressing T cell populations either slowed tumor growth or cleared tumors more efficiently than EV P14 T cells, resulting in significantly improved mouse survival (Figure 1J). Overall, we found that T_PEX_ and T_RM_ shared proteostasis-related gene expression, and overexpression of *Neurl3*, *Rnf149*, or *Wsb1* improved T cell function by maintaining a T_PEX_-like T cell fate in cancer.

### Rescue of T cell function by sustained expression of NEURL3, RNF149 and WSB1 in chronic viral infection

We next tested how ligase overexpression impacted T cell differentiation with LCMV-clone 13, which leads to chronic viral infection^39–41^ (Figure 2A). We found a higher T_PEX_ frequency (CX3CR1^lo^SLAMF6^hi^), decreased T_EX_ frequency (CX3CR1^lo^SLAMF6^lo^) in ligase overexpressing P14 T cells compared to control T cells (Figure 2B). This corresponded with an increased frequency of TCF1^+^ expression in ligase-overexpressing P14 compared to control, but no difference in frequency of TOX^+^ expression (Figure 2C,D). These changes in exhaustion subset frequencies also correspond to increased functionality, with increased frequencies of ligase-overexpressing TNF^+^IFNɣ^+^ P14 T cells after peptide restimulation compared to control P14 T cells (Figure 2E). These data highlight that ligase overexpression favors T_PEX_ accumulation regardless of exhaustion context.

**Figure 2:**
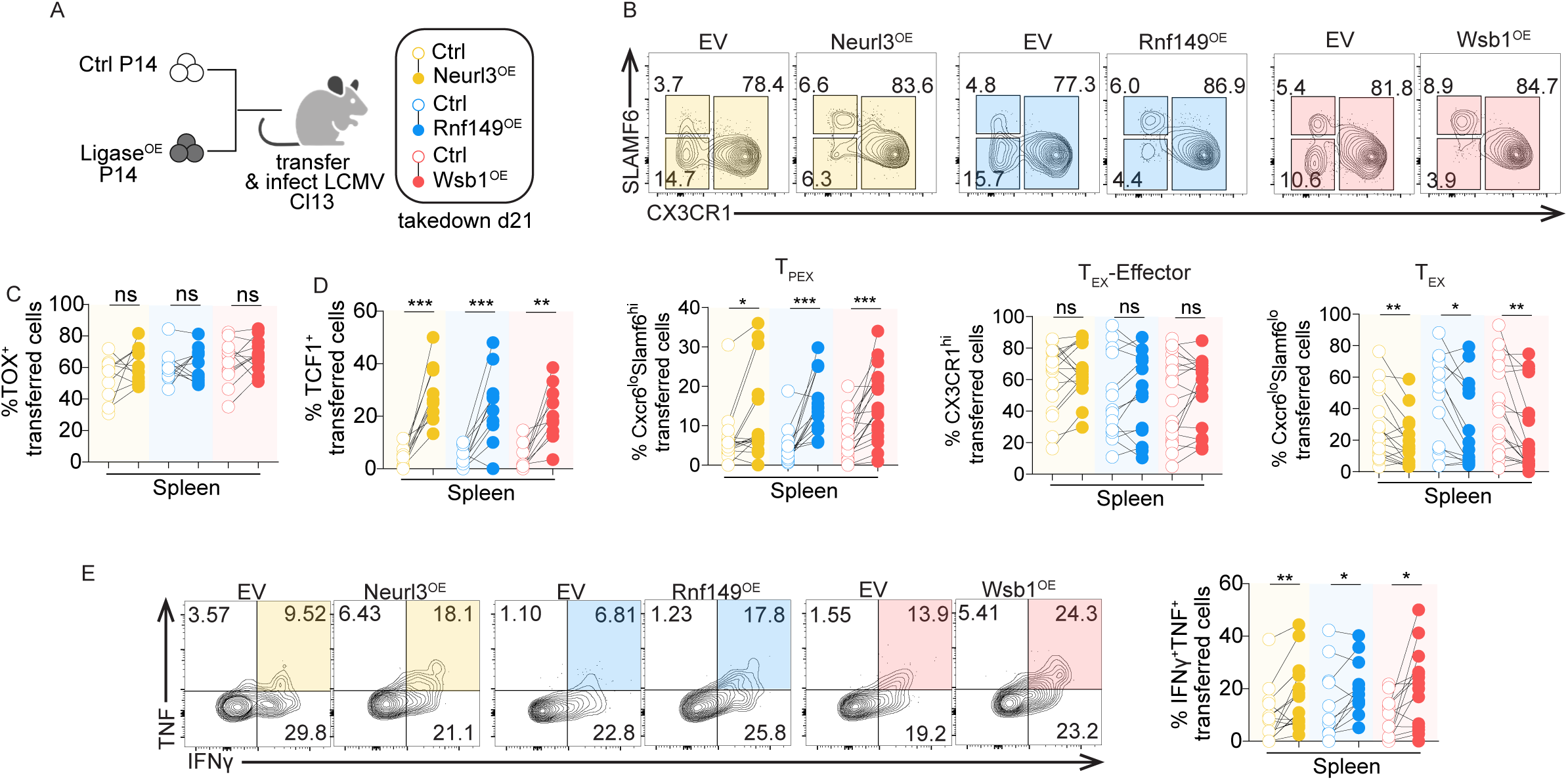
E3 ubiquitin ligases NEURL3, RNF149 and WSB1 expression increases T_PEX_ and decreases T_EX_ frequency in LCMV Cl13 chronic infection. A) Experimental schematic: Congenically mismatched P14 T cells were activated, transduced 24 h later with Empty Vector (EV) or ligase-overexpressing retrovirus (Neurl3^OE^, Rnf149^OE^, or Wsb1^OE^). Next day, cells were sorted on ametrine reporter expression, then EV or Ligase^OE^ cells were mixed 50:50 (1,000 each; 2,000 total) and transferred into recipient mice subsequently infected with LCMV Cl13; spleens were analyzed 21 days later (BioRender). B) Representative flow plots and quantification of SLAMF6 vs CX3CR1 P14 T cell frequencies as in A. Subsets defined as T_PEX_ (CX3CR1^lo^SLAMF6^hi^), T_EX_-Effector (CX3CR1^hi^) and T_EX_ (CX3CR1^lo^SLAMF6^lo^). (*Neurl3*^OE^ n=16; *Rnf149*^OE^ n=14; *Wsb1*^OE^ n=18); five independent experiments. C) Quantification of TOX^+^ P14 T cell frequencies as in A (*Neurl3*^OE^ n=10; *Rnf149*^OE^ n=11; *Wsb1*^OE^ n=12); two independent experiments. D) Quantification of TCF1^+^ P14 T cell frequencies as in A (*Neurl3*^OE^ n=10; *Rnf149*^OE^ n=11; *Wsb1*^OE^ n=10); two independent experiments. E) Representative flow plots and quantification of TNF^+^IFNɣ^+^ P14 T cell frequencies as in A followed by 4 h of peptide restimulation plus protein transport inhibitor (*Neurl3*^OE^ n=15; *Rnf149*^OE^ n=13; *Wsb1*^OE^ n=17); four independent experiments. Statistical significance was calculated using (B-E) paired Student’s *t*-tests. Connecting lines indicate that cells were isolated from the same mouse.

### Loss of NEURL3, RNF149, or WSB1 accelerates T cell exhaustion in tumors and differentiation during acute infection

To understand how loss of NEURL3, RNF149, and WSB1 impacted T cell differentiation, we used CRISPR/Cas9-mediated deletion of each of these ligases and followed the T cell response to both tumor and acute infection. P14 T cells were nucleofected with recombinant Cas9 complexed with trans-activating CRISPR RNA (tracrRNA) and ligase-specific crRNA. Equal numbers of control Thy1^KO^ and ligase^KO^ P14 T cells for each ligase were transferred into B16-GP_33–41_-bearing mice (Figure 3A, Supplemental Figure 3A). Seven days after transfer, all three ligase^KO^ T cell populations were less abundant, had lower TCF1^+^ frequency, and produced less IFN-ɣ upon *ex vivo* restimulation, when compared with their control counterparts in the same host (Figure 3B-E). Taken together, these data indicate that ligase^KO^ T cells become dysfunctional more quickly in comparison to control T cells, with decreased TCF1^+^ frequency and reduced functionality.

**Figure 3:**
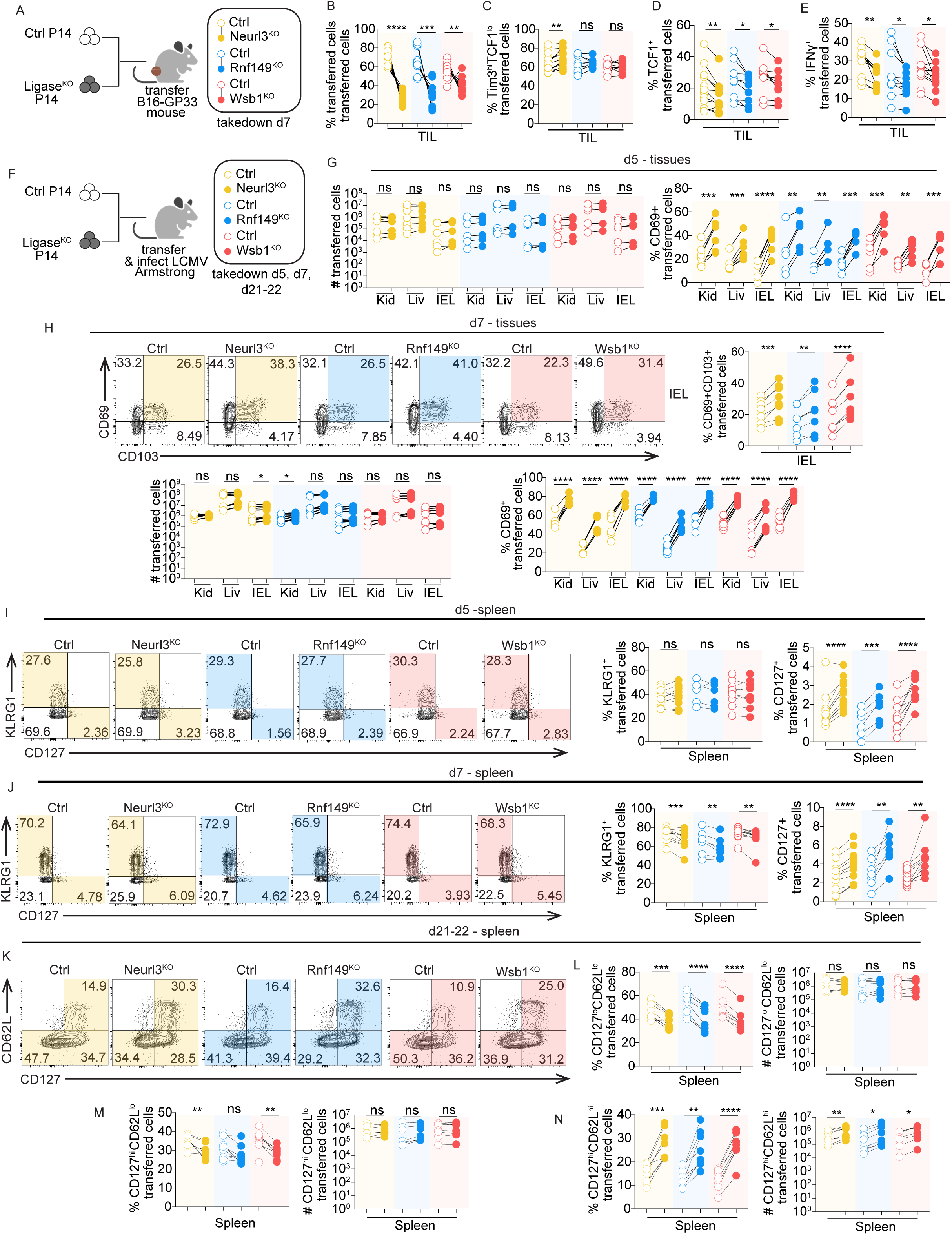
Loss of NEURL3, RNF149, or WSB1 expression accelerates T cell differentiation kinetics. A) Experimental schematic: Congenically mismatched P14 T cells were activated then nucleofected 24 h later with Thy1^KO^ Control CRISPR RNP or ligase^KO^ (Neurl3^KO^, Rnf149^KO^, Wsb1^KO^) plus Thy1^KO^ CRISPR RNP. Cells were cultured to d5-6, mixed 50:50 (1 × 10^6^ each; 2 × 10^6^ total), and transferred into d8 B16-GP_33–41_-bearing mice; dLN and TIL were analyzed 7 days later (BioRender). B) Quantification of transferred P14 T cell frequencies as in A (*Neurl3*^KO^ n=13; *Rnf149*^KO^ n=12; *Wsb1*^KO^ n=12); three independent experiments. C) Quantification of Tim3^hi^TCF1^lo^ P14 T cell frequencies as in A (*Neurl3*^KO^ n=13; *Rnf149*^KO^ n=9; *Wsb1*^KO^ n=9); three independent experiments. D) Quantification of TCF1^+^ P14 T cell frequencies as in A (*Neurl3*^KO^ n=13; *Rnf149*^KO^ n=9; *Wsb1*^KO^ n=9); three independent experiments. E) Quantification of IFNɣ^+^ P14 T cell frequencies as in A followed by 4 h of peptide restimulation plus protein transport inhibitor (*Neurl3*^KO^ n=10; *Rnf149*^KO^ n=11; *Wsb1*^KO^ n=12); three or more independent experiments. F) Experimental schematic: Congenically mismatched P14 T cells were activated and nucleofected 24 h later with Thy1^KO^ Control CRISPR RNP, or ligase KO (*Neurl3*^KO^, *Rnf149*^KO^, *Wsb1*^KO^) plus Thy1^KO^ CRISPR RNP. EV and ligase^KO^ cells were mixed 50:50 (100,000 each; 200,000 total) and transferred into recipient mice subsequently infected with LCMV Armstrong. At 5 or 7 days, circulating CD8^+^ T cells were IV labeled before sacrifice, and tissues were analyzed (BioRender). G) Quantification transferred P14 T cell numbers and CD69^+^ frequencies in the kidney, liver, and IEL at day 5 as in F (*Neurl3*^KO^ n=7; *Rnf149*^KO^ n=6; *Wsb1*^KO^ n=6); two independent experiments. H) (Top left) Representative CD69 vs CD103 flow plot in the IEL; (Top right) CD69^+^CD103^+^ T cell frequencies in the IEL; (Bottom left) transferred P14 T cell numbers; and (Bottom right) CD69^+^ P14 T cell frequencies in the kidney, liver, and IEL at day 7 as in F (*Neurl3*^KO^ n=7; *Rnf149*^KO^ n=7; *Wsb1*^KO^ n=7); two independent experiments. I) (Left) Representative KLRG1 vs CD127 flow plot in the spleen; (Right) Splenic KLRG1^+^ or CD127^+^ P14 T cell frequencies at day 5 as in F (*Neurl3*^KO^ n=11; *Rnf149*^KO^ n=6; *Wsb1*^KO^ n=10); two or more independent experiments. J) (Left) Representative KLRG1 vs CD127 flow plot in the spleen; (Right) splenic KLRG1^+^ or CD127^+^ P14 T cell frequencies at day 7 as in F (*Neurl3*^KO^ n=11; *Rnf149*^KO^ n=7; *Wsb1*^KO^ n=11); two or more independent experiments. K) Representative splenic P14 CD62L vs CD127 flow plots at 21-22 days as in F. L) Frequencies and cell numbers of splenic CD127^lo^CD62L^lo^ P14 T cells at day 21-22 as in F (*Neurl3*^KO^ n=8; *Rnf149*^KO^ n=8; *Wsb1*^KO^ n=8); two independent experiments. M) Frequencies and cell numbers of splenic CD127^hi^CD62L^lo^ P14 T cells at day 21-22 as in F (*Neurl3*^KO^ n=8; *Rnf149*^KO^ n=8; *Wsb1*^KO^ n=8); two independent experiments. N) Frequencies and cell numbers of splenic CD127^hi^CD62L^hi^ P14 T cells at day 21-22 as in F (*Neurl3*^KO^ n=8; *Rnf149*^KO^ n=8; *Wsb1*^KO^ n=8); two independent experiments. Statistical significance was calculated using paired Student’s *t*-tests (B-E, G-N). Connecting lines indicate that cells were isolated from the same mouse.

To test if NEURL3, RNF149, and WSB1 deletion changed T cell differentiation in acute infection, control and ligase^KO^ P14 T cells were transferred into mice that were then infected with LCMV Armstrong, and T cells were isolated on d5, d7, and d21-22 of infection (Figure 3F). While there were no differences in the number of ligase^KO^ T cells compared to control T cells on d5, we observed increased CD69 expression by ligase^KO^ T cells in the tissue on both d5 and d7 post-infection, including more CD69^+^CD103^+^, CD103^+^, and CCR9^+^ ligase^KO^ T cells in the IEL, suggesting that ligase loss alters differentiation kinetics early after acute infection (Figure 3G-H, Supplemental Figure 1B, 3B,C). In the spleen, ligase deficiency resulted in a higher frequency of CD127^+^ memory-precursor T cells on d5 of infection, but there was no difference in the frequency of KLRG1^+^ short-lived effector T cells (Figure 3I). By d7 post-infection, we observed a decline in the frequency of KLRG1^+^ T cells, which suggested that the ligase^KO^ T cells were already beginning to contract (Figure 3J). At memory timepoints (days 21–22 post-infection), we found that higher frequencies of CD69^+^ ligase^KO^ T cells were maintained in the tissues, and that there was an increase in the number of ligase^KO^ T_RM_ in IEL (Supplemental Figure 3D). In the spleen, we observed decreased frequencies of long-lived, terminal-effector (CD62L^lo^CD127^lo^) and effector-memory (CD62L^lo^CD127^hi^) T cell subsets, with a corresponding increase in the central-memory (CD62L^hi^CD127^hi^) T cell subset, revealed both by frequency and absolute cell number of ligase^KO^ T cells compared with the control T cells, indicating that deficiency in all three ligases had a long-term impact on cell fate (Figure 3K-N). Finally, to directly interrogate the differentiation potential of ligase-deficient T cells, we adoptively transferred control or ligase^KO^ P14 cells into mice that were then infected with LCMV Armstrong. On day 30 of infection, T_CM_ from control or ligase^KO^ P14 T cells were sorted and then co-transferred at a 50:50 ratio into new recipients followed by LCMV Armstrong infection to assess secondary memory differentiation potential. This experiment showed that ligase-deficient T cells were less abundant than control T cells at secondary memory timepoints across tissue compartments (Supplemental Figure 3E). Thus, we found that in response to tumors and to acute infection, loss of ligase expression led to altered T cell differentiation in circulating memory, tissue-resident memory, and TIL populations.

To characterize and compare the functional impact of NEURL3, RNF149, and WSB1 on the T cell transcriptome, we performed scRNA-seq analysis of control and ligase^KO^ P14 T cells on d5 of infection with LCMV Armstrong, when we observed early phenotypic differences. We found that ligase deficiency in either the spleen or IEL shared RNA scRNA-seq clusters with the control and analysis revealed few differentially expressed genes enriched in the ligase^KO^ T cells, indicating that ligase loss-of-function minimally altered the transcriptome. (Supplemental Figure 4A,B, Supplemental Table 2). Consistent with this observation, RNA-seq performed on EV and Neurl3^OE^ P14 T cells obtained from MC38-GP_33–41_ tumors, revealed few significant differences in gene expression between EV and Neurl3^OE^ T cells in the dLN or tumor (Supplemental Figure 4C,D). Taken together, these data indicate that NEURL3, RNF149, and WSB1 impact T cell differentiation, but the presence or absence of these ligases minimally impacted mRNA expression. E3 ligases have been previously shown to control self-renewal and cell-fate determination in intestinal stem cells, hematopoietic stem cells, and embryonic development through targeted protein degradation^15,42^, and therefore NEURL3, RNF149, and WSB1 may be having a similar function in CD8^+^ T cell differentiation.

### Comparing proteostasis in *ex vivo* T_EX_, T_PEX_, T_RM_, and T_EFF_

As NEURL3, RNF149, and WSB1 likely influence cell differentiation potential and cell fates via targeted protein degradation, we used quantitative proteomics to measure protein abundance in T cells. The extent to which TIL mRNA corresponds to protein expression has mainly been characterized on a gene-by-gene basis and relative mRNA versus protein comparisons have been made for *in vitro*-cultured T cells, lymphocytes at steady-state, and other non-immune cell types, but similar evaluations in *ex vivo* T cells upon infection or tumor formation have been undercharacterized^23,27,43–46^. To understand the proteome in *ex vivo* T cell populations, we used mass spectrometry to compare proteomes in T_EX_ and T_PEX_ from tumors and T_RM_ and T_EFF_ from acute infection (Figure 4A). We isolated P14 T cells from the spleen at the peak of immune response (T_EFF_) and memory timepoint from IEL (T_RM_) following LCMV Armstrong infection, as well as endogenous TIL (T_PEX_ and T_EX_) from B16-bearing mice. We performed scRNA CITE-seq on these populations, as well as proteomic analysis using low-input data-independent acquisition (DIA) mass spectrometry and obtained >8,000 unique proteins per replicate from 1 x 10^5^ sorted cells (Figure 4B,C, Supplemental Tables 3,4). We used TotalSeq oligonucleotide-conjugated antibodies to measure PD-1 and TIM-3 protein expression on TIL in our scRNA-seq data and compared gene-signature enrichment with previously published datasets^3,10^, confirming that the T cell populations identified by scRNA-seq were comparable to the bulk sorted T cells analyzed by mass spectrometry (Supplemental Figure 4E,F). To ensure sufficient cell numbers for analysis, we included scRNA-seq data for T_RM_ from both endogenous CD8⁺ T cells and P14 T cells (Supplemental Figure 4G). Direct comparison of RNA and protein abundances for individual genes revealed that for some genes, RNA and protein showed a linear relationship (quadrant 1 [Q_1_], [Q_3_]), while other genes had greater protein abundance than RNA expression [Q_2_], or greater RNA expression than protein abundance [Q_4_] (Figure 4D, Supplemental Figure 4H). Performing pathway enrichment to identify the top three Gene Ontology Biological Processes (GO:BP) pathways in each quadrant revealed that genes with greater protein than RNA expression were enriched for metabolite pathways, while genes with greater RNA than protein expression included enrichment in cytokine pathways, consistent with observations that T_RM_ and other effector T cell populations can maintain cytokine mRNA in a translationally repressed state^47,48^ (Figure 4E). Using a distance matrix to summarize the relationships between populations categorized by RNA or protein abundance, we observed different correlation patterns: proteomic analyses revealed that T_PEX_ and T_EX_ were more distinct from each other than suggested by transcriptomic data alone (Figure 4F), highlighting how integrating proteomic analyses of *ex vivo* T cells with RNA expression can improve our understanding of T cell biology.

**Figure 4:**
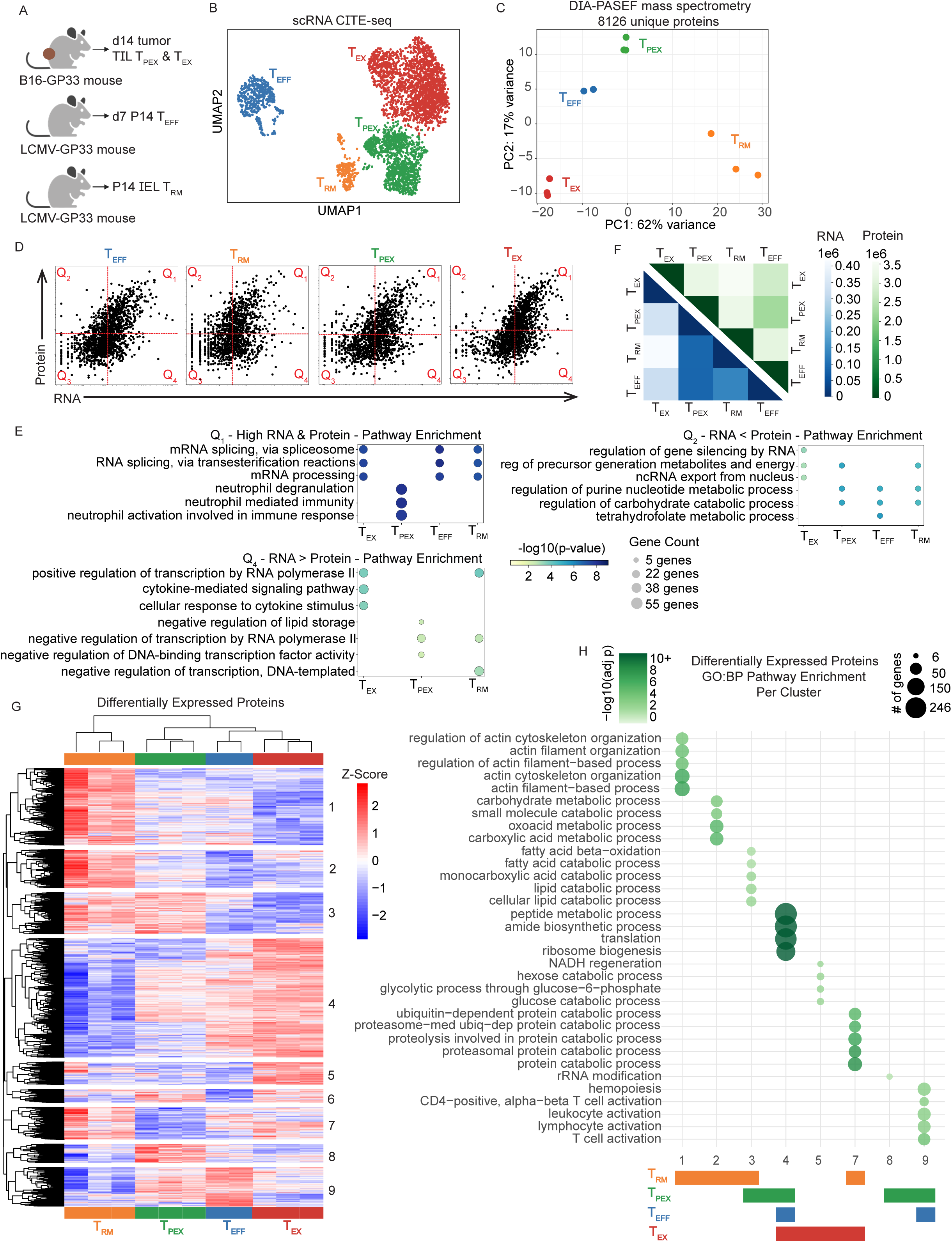
Comparing *ex vivo* T_EX_, T_PEX_, T_RM_, and T_EFF_ transcriptome and proteome. A) Experimental schematic: Endogenous B16 TIL were sorted at d14 (T_PEX_ sorted on CD8^+^PD1^lo/int^Tim3^−^ and T_EX_ sorted on CD8^+^PD1^hi^Tim3^+^). In parallel, P14 were sorted from d7 LCMV Armstrong infection from spleen (T_EFF_), and at memory timepoint from the IEL (T_RM_) (BioRender). B) RNA UMAP from CITE-seq of CD8^+^ T cells from d7 LCMV Armstrong spleen (P14), d30 IEL (P14 + endogenous), and endogenous B16 TIL T_PEX_ and T_EX_ (defined by PD1 vs Tim3 CITE-seq and enrichment signature; Supplemental Figure 4E). The sequenced cells were derived from 1 mouse for d7 spleen, pooled from 6 mice for d30 IEL, and pooled from 10 mice for TIL. C) Protein PCA (DIA-PASEF) of CD8^+^ T cells from d7 LCMV Armstrong spleen (P14), d14 IEL (P14), and endogenous B16 TIL T_PEX_ and T_EX_. D14 T_RM_ was used as an early memory timepoint due to cell number limitations at d30. Replicates: d7 spleen n=2, d14 IEL n=3 (6 pooled samples/replicate), T_PEX_ n=3 (6 pooled/replicate) and T_EX_ n=3 (6 pooled/replicate). D) RNA-protein scatter per T cell subset (non-zero RNA vs non-zero protein, one dot per gene), split into four quadrants by mean expression of total *x*- or *y*-axis. E) Dot plot of genes from either Q_1_, Q_2_, or Q_4_ for each T cell subset (from D), with GO:BP pathway enrichment for the top 3 significantly pathways per subset. F) Euclidean distance heatmap of pseudobulk RNA-seq samples (in blue) and protein intensities (in green) in PCA space (darker = more similar). G) Heatmap with unsupervised hierarchical clustering of the z-score of significant differentially expressed proteins from T_EFF_, T_RM_, T_PEX_, and T_EX_, with adjusted p-value <0.05. H) Gene Ontology bubble plot of proteins defining each cluster from G.

Analysis of differentially expressed proteins revealed nine unique clusters (Figure 4G, Supplemental Table 5). Performing gene ontology pathway analysis of the top five pathways per cluster revealed unique biological processes associated with each cluster (Figure 4H). We found that T_RM_ had increased levels of proteins associated with roles in actin organization and the catabolism of small molecule pathways (clusters 1 and 2) and shared upregulation of proteins related to lipid catabolic processes with T_PEX_ (cluster 3). T_RM_-specific expression of actin-regulatory proteins may indicate a poised activation state, as the immunological synapse requires dynamic actin regulation^49^ or it may be related to the motile nature of T_RM_, as surveying tissue requires dynamic actin regulation for movement^50,51^. For lipid metabolic pathways, lipid catabolism has been previously associated with tissue adaptation and function in T_RM_, and with improved T cell function in cancer^11,52^. We found that T_PEX_ and T_EFF_ both shared proteins with roles in T cell differentiation (cluster 9), and T_EX_ uniquely upregulated proteins related to glycolysis (cluster 5), consistent with previous reports that T_EX_ upregulate glycolytic activities^53^. We found T_EX_ shared the upregulation of proteasome-mediated, ubiquitin-dependent protein catabolism pathways with T_RM_ (cluster 7), an aspect of T_EX_ biology that had not yet been characterized. Similar analysis of differential gene expression followed by GO BP pathway enrichment using RNA revealed the top RNA-enriched pathways were distinct from most pathways observed in the proteomic data (Supplemental Figure 5). Thus, proteomic analyses revealed distinct enrichment of proteasome-mediated protein catabolism pathways in T_EX_ and T_RM_ and uncovered unique proteomic profiles for each of the T cell populations.

### T_EX_ have functional proteasomes, yet accumulate short-lived proteins

Performing GSVA on differentially expressed proteins, we found that both T_EX_ and T_RM_ had increased expression of proteins involved in a number of pathways related to proteasome-dependent degradation relative to T_EFF_ and T_PEX_ (a pattern not reflected in GSVA of differentially-expressed RNA), including protein K48-linked ubiquitination (Figure 5A, Supplemental Figure 6A). We cross-validated our DIA-proteomics results with tandem mass tag (TMT)-based mass spectrometry on both related experimental groups and additional populations: P14 from dLN and B16-GP_33–41_ TIL to allow us to compare endogenous to antigen-specific T cells, T_EM_ and T_RM_ P14 post-LCMV Armstrong infection to compare multiple memory T cell populations, and endogenous T_PEX_ and T_EX_ TIL from B16-GP_33–41_ to validate our DIA-proteomic T_PEX_ and T_EX_ populations (Supplemental Figure 6B,C, Supplemental Table 6). While fewer proteins were identified compared to the DIA-PASEF dataset, we observed similar patterns of protein expression (Supplemental Figure 6D). Using GSVA for differentially expressed proteins followed by GO:BP pathway analysis, we saw similar enrichment in protein degradation and K48-linked ubiquitination in T_EX_ and T_RM_ compared to T_PEX_, but also found that antigen-specific dLN and TIL P14 had similar relative abundance of the pathway proteins to T_EX_, indicating that antigen-specific TIL and endogenous T_EX_ showed a similar phenotype (Supplemental Figure 6E). We also found T_EM_ had similar protein degradation pathway phenotype to T_PEX_ rather than T_RM_, indicating that not all memory T cells upregulate proteasomal degradation and K48-linked ubiquitination equivalently (Supplemental Figure 6E).

**Figure 5:**
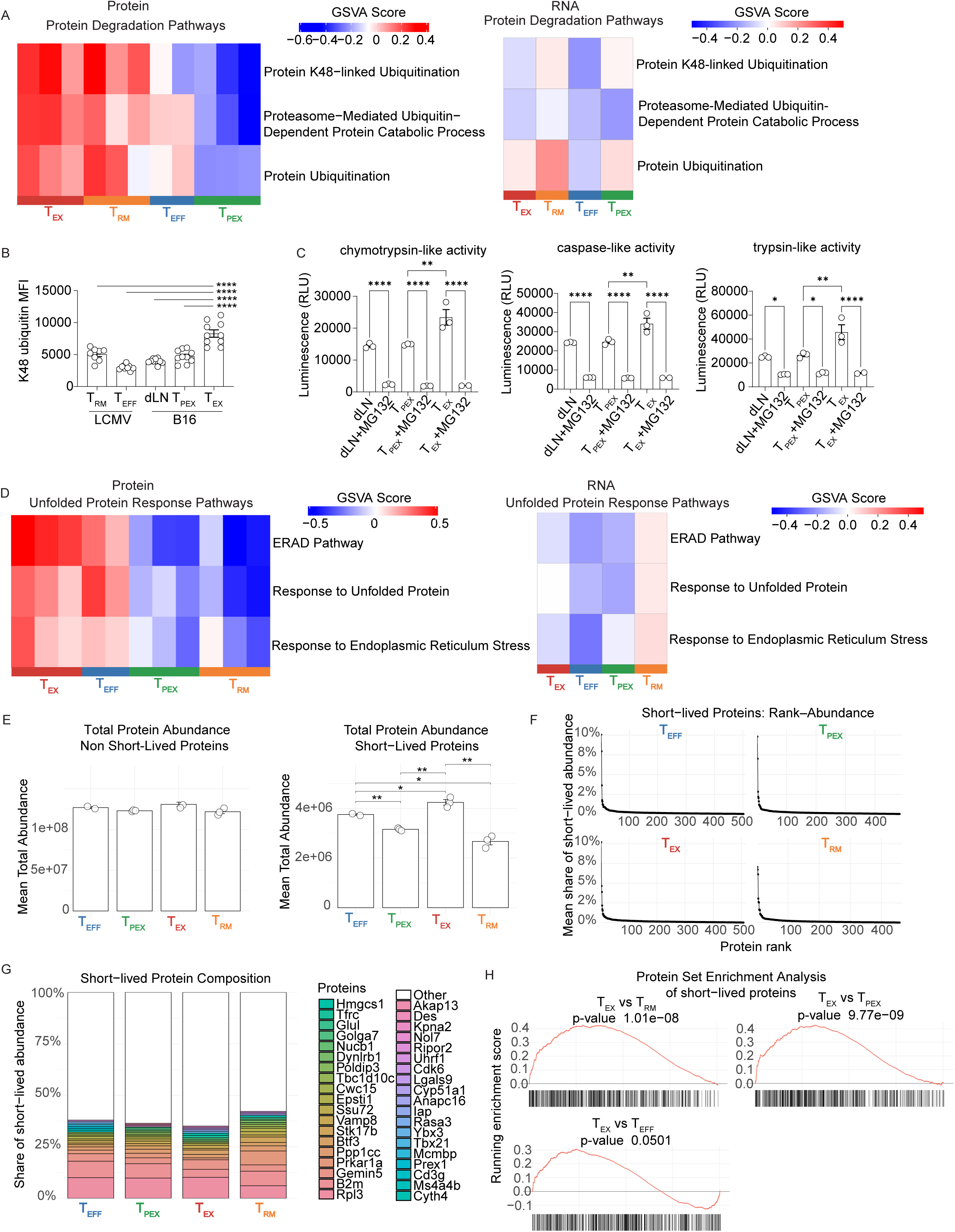
T_EX_ accumulates short-lived proteins yet have functional proteasomes. A) GSVA heatmaps of protein-degradation GO:BP pathways from proteomics (left, data from 4C) and mRNA (right, data from 4B). B) K48 ubiquitin MFI in CD8^+^ T cells from P14 IEL d30 (n=8) and P14 spleen d7 (n=9) after LCMV Armstrong infection; and endogenous dLN (n=10), T_PEX_ (n=10) and T_EX_ (n=10) from B16 melanoma after treatment with 10 µM MG-132 for 1 h at 37°C prior to fixation and staining. C) Chymotrypsin-, caspase-, and trypsin-like activities in sorted dLN, T_PEX_, and T_EX_ incubated ± 10 µM proteasome inhibitor MG-132 (1 h, 37°C, inhibitor as negative control); 2–3 independent replicates. D) GSVA heatmaps of unfolded protein response GO:BP pathways from proteomics (left, data from 4C) and mRNA (right, data from 4B). E) Using *in vitro* T cell protein half-life data (Supplemental Table 7), the *ex vivo* T_EFF_, T_RM_, T_PEX_, and T_EX_ proteomics data were stratified into non-short-lived proteins (half-life between >8 h and ≤20 h; Left) and short-lived proteins (half-life ≤8 h; Right). The mean total abundance per category were quantified. F) For short-lived proteins (defined in E), rank-abundance curves by subset. G) For short-lived proteins (defined in E), composition shown as 100% stacked bars with the top 20 proteins per cell type and remaining proteins collapsed into “other”. H) For short-lived proteins (defined in E), Protein Set Enrichment Analysis across pairwise comparisons with T_EX_, plots show running enrichment score and short-lived protein postion. Statistical significance was calculated using (B, C, E) one-way ANOVA and empirical Bayes–moderated t-statistic (G).

To compare the proteostasis status of T_RM_ to additional circulating memory T cell populations, we performed DIA-PASEF mass spectroscopy on P14 T cells sorted on day 33 post LCMV-Armstrong infection from the spleen (CD127^hi^CD62L^hi^ T_CM_, CD127^hi^CD62L^lo^ T_EM_, CD127^lo^CD62L^lo^ LLTE) and the IEL (T_RM_) (Supplemental Figure 7A). We obtained >6,000 unique proteins per replicate from 2 x 10^3^ sorted cells (Supplemental Figure 7B) and integrated these data with results from Figure 4C by normalizing the datasets to their shared T_RM_ population (Supplemental Figure 7C). We then compared these data to our previously identified cell-type specific protein expression from Figure 4G and found that all three circulating memory T cell populations shared substantial protein expression with T_RM_-enriched proteins but did not share protein expression with T_PEX_+T_RM_, T_EFF_+T_PEX_+T_EX_, or the T_EX_ protein clusters (Supplemental Figure 7D). Using GSVA for differentially-expressed proteins followed by GO:BP pathway analysis on the combined datasets, we observed similar enrichment in protein degradation and K48-linked ubiquitination in T_EX_ and T_RM_ compared to T_PEX_, T_EFF_, T_CM_, T_EM_, and LLTE (Supplemental Figure 7E).

Increased abundance of K48-linked polyubiquitin chains can be used to indicate defective protein degradation, therefore we measured K48 ubiquitin staining by flow cytometry and found significantly increased levels of K48-linked polyubiquitin in T_EX_ and T_RM_; however, K48-linked polyubiquitin was higher in T_EX_ compared to other CD8^+^ T cell groups, suggesting that T_EX_ may have increased proteostasis dysfunction compared to other T cell populations (Figure 5B). To determine if proteasome-mediated protein degradation was functional in the T_EX_ population, we utilized proteasome peptidase activity assays and found that T_EX_ possessed functional proteasomes, as assessed by increased chymotrypsin-like, caspase-like, and trypsin-like proteasome activities compared with the dLN or T_PEX_ populations (Figure 5C).

Interestingly, GO pathway analysis revealed that both T_EX_ and T_EFF_ had higher levels of proteins involved in UPR pathways, a pattern not reflected in RNA expression (Figure 5D, Supplemental Figure 6A). In our TMT-mass spectrometry dataset, we saw both endogenous T_EX_ and antigen-specific P14^+^ TIL had similarly upregulated proteins involved in the UPR pathways, while endogenous T_PEX_, P14 T_EM_, and P14 T_RM_ had decreased relative abundance compared to T_EX_ (Supplemental Figure 6F). In comparison to circulating memory T cells, we also observed enrichment in response to unfolded protein and ER stress response (hallmarks of UPR) only in T_EX_ and T_EFF_, but not in T_RM_, T_PEX_, T_CM_, T_EM_, and LLTE (Supplemental Figure 7F). The UPR in TIL has previously been characterized as negatively correlated with T cell function in tumors, since studies limiting UPR activation via IRE1α–XBP1 or PERK–CHOP showed improvement of TIL function^54–57^. To understand proteostasis dysregulation and evaluate protein turnover dynamics within TIL, we performed metabolic protein labeling using heavy isotopes of arginine (^13^C_6_, ^15^N_4_) and lysine (^13^C_6_, ^15^N_2_) in *in vitro*-cultured CD8^+^ T cells. This approach allowed us to identify rapidly degraded proteins in mouse CD8^+^ T cells (i.e., half-life ≤8 h, Supplemental Table 7). We then used this data to assign our *ex vivo* proteomics data to short-lived (half-life ≤8 h) and non-short-lived (half-life >8 h to ≤20 h) protein subsets (Figure 5E-H). The abundance of non-short-lived proteins was similar among T_RM_, T_EFF_, T_PEX_, and T_EX_. However, when quantifying the abundance of short-lived proteins, we observed that T_EX_ had an enrichment for numerous short-lived proteins, despite having functional proteasomes (Figure 5E-H). These data indicate that protein degradation in T_EX_ is dysfunctional either due to a proteasome-independent mechanism or that T_EX_ proteasomal subunit composition may be suboptimal for protein degradation *in vivo* (Figure 5C,E, Supplemental Figure 7G).

### Unfolded protein accumulation in T_EX_ is rescued by NEURL3, RNF149, or WSB1 expression

To characterize the conformational states of the proteins in T cells, we stained for unfolded proteins using Proteostat, a dye that fluoresces upon binding to unfolded protein quaternary structures^58^. We found that T_EX_ had significantly increased levels of unfolded proteins compared to T_PEX_, T_RM_, and T_EFF_ cells (Figure 6A,B). Inhibiting protein synthesis *ex vivo* for 6 hr did not change unfolded protein accumulation in T_EX_, despite observing a decrease in Cyclin A2 (a rapidly degraded protein^59^) during this timeframe (Figure 6C). These data indicate that the increased levels of unfolded proteins in T_EX_ do not primarily arise from newly synthesized proteins or from dysfunctional proteasomes.

**Figure 6:**
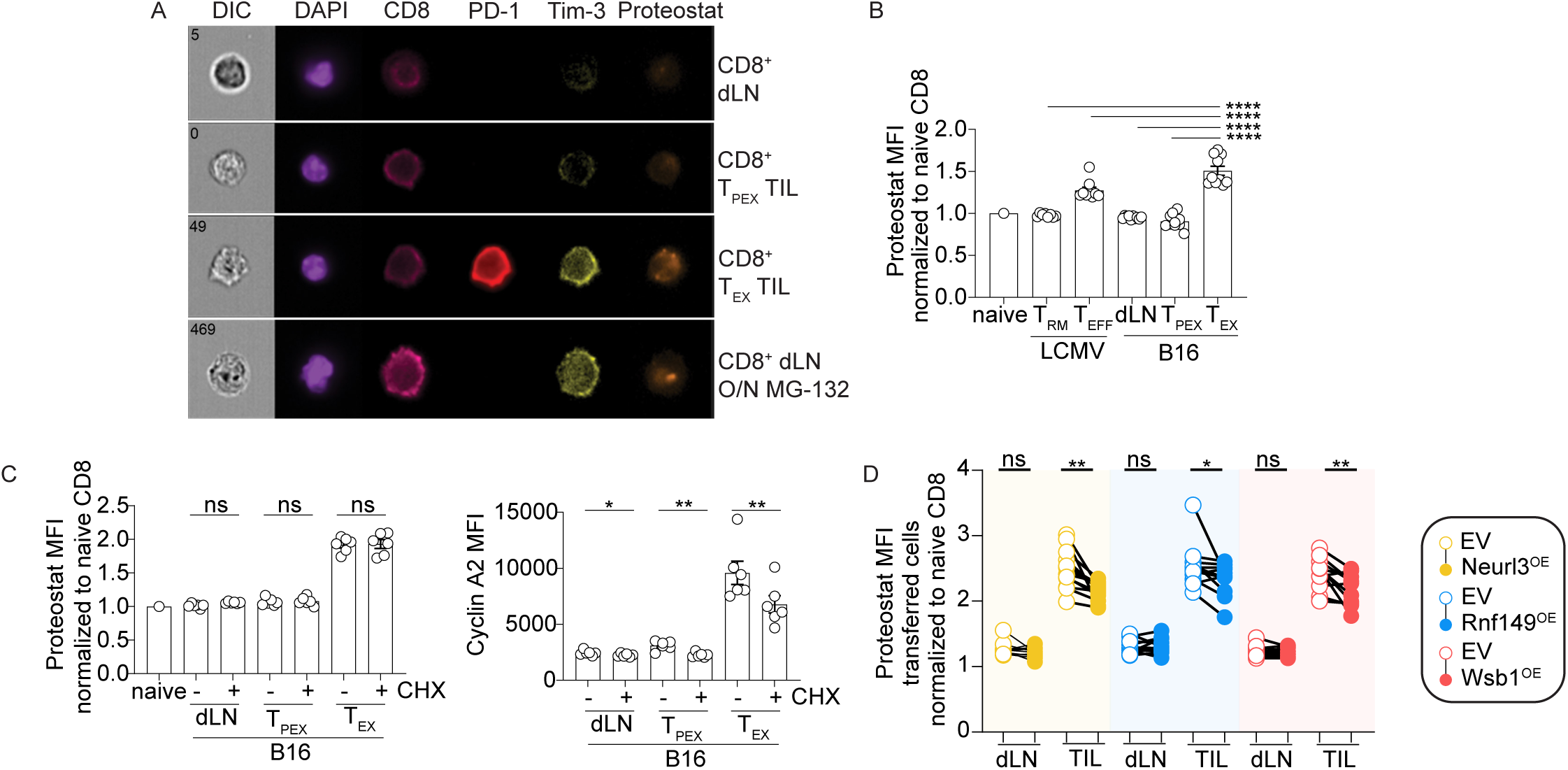
T_EX_ accumulates unfolded proteins, rescued by NEURL3, RNF149, or WSB1 expression. A) ImageStream visualization of Proteostat staining of CD8^+^ T cells isolated from dLN or TIL (T_PEX_ and T_EX_), plus MC-132-treated CD8^+^ T cells (5 µM, 16hr). B) Proteostat MFI (normalized to spiked-in, congenically mismatched naive CD8^+^ T cells) in P14 IEL d30 (n=8) or P14 spleen d7 (n=9) after LCMV Armstrong infection, and endogenous dLN (n=10), T_PEX_ (n=10) and T_EX_ (n=10) TIL from B16 melanoma. C) (Left) Proteostat MFI (normalized as in B) and (Right) Cyclin A2 MFI from CD8^+^ T cells isolated from endogenous dLN (n=6), T_PEX_ (n=6) and T_EX_ (n=6) TIL from B16 melanoma, treated ± Cycloheximide (20 µg/mL, 6 h, 37°C). Representative of three independent experiments. D) Proteostat MFI (normalized as in B) from P14 T cells from dLN or tumors of mice with congenically distinct EV vs *Neurl3*^OE^, *Rnf149*^OE^, or *Wsb1*^OE^ 8 days after adoptive transfer into tumor-bearing mice (*Neurl3*^OE^ n=12; *Rnf149*^OE^ n=12; *Wsb1*^OE^ n=11); three independent experiments. Statistical significance was calculated using (B) one-way ANOVA and paired Student’s *t*-tests (C-D). Connecting lines indicate that cells were isolated from the same mouse.

Since T_EX_ proteasomes were functional and protein synthesis did not drive unfolded protein accumulation, we instead targeted protein degradation upstream of proteasome activity by overexpressing the E3 ubiquitin ligases NEURL3, RNF149, or WSB1 in P14 TIL. We found that there was a minimal impact on dLN T cells, whereas ligase overexpression rescued the elevated unfolded protein accumulation in TIL (Figure 6D). Taken together, these data show that T_EX_ experience a unique proteostasis state in which protein degradation can occur and the UPR is upregulated, yet T_EX_ have significant accumulation of unfolded proteins. Overexpression of NEURL3, RNF149, or WSB1 in T cells rescues accumulation of unfolded proteins, indicating that unfolded proteins contribute to T_EX_ dysfunction and that modulation of specific E3 ubiquitin ligase activity reduces proteotoxic stress.

### NEURL3, RNF149, or WSB1 improve immunotherapeutic responses in humans and improve cancer immunotherapy in mice

We next wanted to understand the therapeutic potential of NEURL3, RNF149, and WSB1 expression in the context of anti-tumor immunity. Quantification of the scRNA-seq data from melanoma patient TIL^60^ showed that *RNF149* and *WSB1* were expressed at higher levels in T_RM_-like TIL and T_PEX_ than in other CD8^+^ T cell populations in the tumor (Figure 7A), consistent with our murine TIL data (Figure 1B). Quantification of *RNF149* and *WSB1* transcripts in patients that received checkpoint blockade showed that prior to the start of therapy, the expression levels of *RNF149* and *WSB1* were moderately predictive of response to therapy, while the levels of *RNF149* and *WSB1* expression following checkpoint blockade were highly correlated with the favorable response to checkpoint blockade (Figure 7B). Human scRNA-seq analysis did not detect the *NEURL3* transcript, however, bulk RNA-seq from patients with metastatic melanoma who had their primary tumor resected and sequenced prior to checkpoint blockade immunotherapy^61^ showed higher levels of *NEURL3*, *RNF149*, or *WSB1* expression in patients that had improved survival following checkpoint blockade immunotherapy (Figure 7C). Thus, ligase expression correlates with improved responses to immunotherapy in humans.

**Figure 7:**
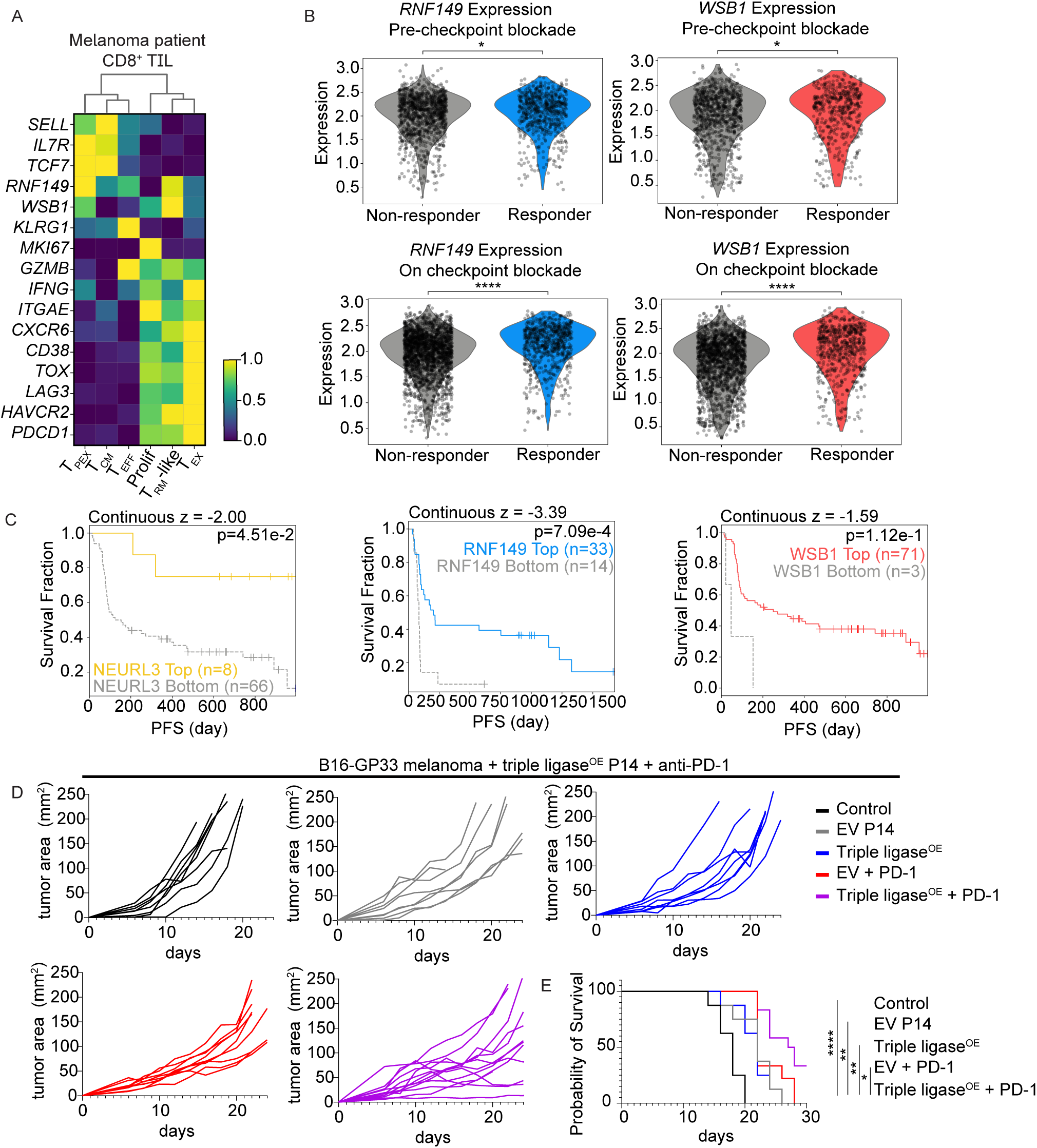
NEURL3, RNF149, or WSB1 expression corresponds with immunotherapeutic responses to checkpoint blockade in human melanoma and improves cancer immunotherapy in mice. A) Heatmap of CD8^+^ T cell clusters from checkpoint blockade-treated melanoma patients (GSE120575^60^), annotated by top gene expression per cluster; RNF149 and WSB1 expression shown. B) Violin plots of RNF149 or WSB1 RNA expression in CD8^+^ TIL with patient data from A, stratified into non-responder vs responder, pre-treatment (top) and on-treatment (bottom). C) Survival of patients with melanoma (dbGaP Study Accession: phs000452.v3.p1^61^) treated with checkpoint blockade after primary tumor resection and sequencing, stratified by gene expression (high/top vs low/bottom) in the primary tumor. The survival risk score is *z*-score gene effect of death risk in a Cox proportional hazards model. Analysis conducted by TIDE^79^. D) B16-GP_33–41_ tumor area in mice that received no T cells, EV P14 or *Neurl3*^OE+^*Rnf149*^OE+^*Wsb1*^OE^P14 T cells on d6 after tumor cell implantation with isotype or anti-PD1 treatment every other day; two independent experiments. E) Survival from D. Statistical significance was calculated using the (B) unpaired Student’s *t*-tests or (C) CoxPH test or the (E) log-rank (Mantel-Cox) test.

To determine if ligase overexpression could improve T cell responses to B16-GP_33–41_ mouse melanoma in a tumor resistant to checkpoint blockade, we combined cell therapy and anti-PD1 antibody treatment. P14 T cells were transduced with all three ligases and adoptively transferred into mice with established B16-GP_33–41_ melanoma and treated with either isotype control or anti-PD1 therapy. We found that triple ligase^OE^ cell therapy with checkpoint blockade improved tumor control and significantly improved survival compared to EV P14 and triple ligase^OE^ isotype treated mice (Figure 7D,E). Thus, ligase-expressing CD8^+^ T cell therapy synergizes with checkpoint blockade and improves immunotherapeutic responses to cancer, highlighting the role of proteostasis in regulating anti-tumor T cell responses during immunotherapy targeting co-inhibitory receptors.

## Discussion

In this study, we sought to understand the relationship between T cell function in tumors versus tissues given the association of T_RM_-like TIL and improved responses to immunotherapy in patients with cancer^9^. The identification of T_RM_-associated genes lost with terminal differentiation to T_EX_ reveals new avenues for understanding exhaustion, with downregulation of protein folding and degradation gene-expression profiles suggesting that T_EX_ experience disrupted proteostasis, while T_PEX_ and T_RM_ actively maintain proteostasis. Active maintenance of differentiation potential by sustained proteostasis may be an important biological difference between memory and naive T cells: proteasome activity has been shown to be crucial for generating circulating memory T cell populations, and memory T cells maintain a higher level of protein turnover than naive T cells prior to TCR stimulation, while quiescent naive T cells and quiescent stem cells both have low levels of protein synthesis and degradation^27,28,62,63^. Previous research supports the roles of NEURL3, RNF149, and WSB1 in maintaining differentiation potential and responding to cell stress in non-immune cells, which may also help to maintain differentiation potential^64^. NEURL3 was previously shown to target Vimentin for degradation, to inhibit the epithelial mesenchymal transition of nasopharyngeal carcinoma^65^. RNF149 targets BRAF for degradation in cancer cells, reduces reactive oxygen species in cell lines, and regulates protein quality control in the ER during cell stress^66–68^. WSB1 was observed to be expressed under hypoxic conditions, and has been reported to target VHL, DNA damage-responsive ataxia-telangiectasia mutated (ATM) serine/threonine kinase, and Rho-binding protein RhoGDI2 for degradation^69–71^. WSB1 also correlates with increased tumor cell metastasis, although it exerts a protective effect against Parkinson’s disease by targeting the leucine-rich repeat kinase 2 protein (LRRK2) in neurons^69,71–73^. It is not clear from our work what the specific targets of NEURL3, RNF149, or WSB1 are in T cells or if targets are shared across cell types. Given that these ligases do not share sequence homology, each ligase may target unique proteins, converging on a similar stem-like phenotype. This may include proteins in the TCR signaling cascade, proteins involved in RNA transcription, or proteins involved in protein translation, or additional pathways that help enforce a stem-like state. Recent publications identifying E3 ligase targets in cell lines using proximity labeling showed many E3 ligases can have hundreds of protein targets^74,75^, rather than one predominant target. Defining the protein targets for NEURL3, RNF149, and WSB1 will be an important topic for future work.

Mass spectrometry of immune cells has been used as a tool to reveal metabolic states of *in vitro* and *ex vivo*-isolated T cells^76^, but it has previously required large cell numbers for the acquisition of in-depth protein information, making *ex vivo* assessment of rare populations technically challenging. DIA-PASEF mass spectroscopy and TMT mass spectrometry^45^ are important technological advancements, allowing for proteomic assessments of less-abundant populations and yielding higher sensitivity and deeper proteome coverage. Our data is consistent with reports that mRNA profiles poorly correlate with protein expression patterns^23,27,44–46^, highlighting that comparing mRNA and protein provides complementary yet unique biological information. This may be a particularly important insight for T_EX_, as the RNA profiles in T_EX_ do not clearly explain why T_EX_ are dysfunctional or how checkpoint blockade permits some patients to respond to therapy while others do not. Proteomics may help reveal information about T_EX_ and other immune cell populations to aid immunotherapy development.

Our mass spectrometry datasets have revealed shared pathway enrichment in T_RM_ and T_EX_ proteasomal degradation, yet only T_EX_ upregulate UPR, despite having functional proteasomes (Figure 5C,D). Complementary findings examining disrupted proteostasis by T_EX_ were recently reported, further characterizing T_EX_ as not only having unfolded proteins, but protein aggregates and stress granules as well^46^. Comparing T_EX_ to cells from other unfolded protein diseases, such as neurodegenerative diseases, may help us to understand the biological consequences of unfolded protein accumulation in T_EX_^20^: unfolded proteins themselves or ER stress driven by genetic or environmental factors can trigger the UPR, causing cells to respond transcriptionally to the proteotoxic stress. If this response is successful, the cell will return to proteostasis, known as “adaptive UPR”^20^. Conversely, if the UPR is unable to resolve the proteotoxic stress, this “terminal UPR” state results in cell dysfunction^20^. From our data, we see that T_EX_ upregulate the UPR and yet still have an abundance of unfolded proteins, suggesting that T_EX_ also experience a terminal UPR state. It is unclear whether NEURL3, RNF149, or WSB1 directly reduce the accumulation of unfolded proteins resulting from targeted degradation or instead prevent this phenotype through indirect mechanisms, but since all three ligases rescue unfolded protein accumulation, it suggests that this phenotype is a broader consequence of disrupted proteostasis involving multiple proteins rather than the dysfunction of a single unfolded protein. Proteotoxic cell stress has been shown to be associated with resistance to checkpoint blockade in human cancers^77^, therefore engineering T_EX_-specific proteolysis-targeting chimeras (PROTACs), or ligase overexpression in chimeric antigen receptor (CAR) T cell or other cellular immunotherapies, could be used for immunotherapy in human cancer.

## Limitations of the study

We identify NEURL3, RNF149, and WSB1 as molecular contributors to the maintenance of CD8^+^ T cells differentiation potential in both cancer and chronic viral infection, but the direct targets remain undefined. Proximity-labeling studies of other E3 ubiquitin ligases have identified hundreds of potential targets, suggesting that these ligases may similarly act through a broad network. It also remains unclear whether unfolded protein accumulation in T_EX_ is a cause or a consequence of dysfunction, therefore future studies will be required to clarify the role of unfolded protein in the overall biological state of T cell exhaustion. In addition, the analysis of human T cell exhaustion is currently limited to gene expression in melanoma. Extending these findings to other human cancer types as well as proteome profiling will be essential to assess the human TIL proteostasis state.

## Resource availability

### Lead Contact

Requests for resources and reagents should be directed to and will be fulfilled by the Lead Contact, Ananda Goldrath.

### Materials availability

Unique reagents generated in this study are available from the lead contact with a completed materials transfer agreement.

### Data and Code Availability

The RNA datasets generated during this study are available at NCBI GEO repository as indicated in the key resources table. The mass spectrometry proteomics datasets generated during this study are available at the ProteomeXchange Consortium via the PRIDE partner repository or MassIVE as indicated in the key resources table. This paper does not report original code.

## Acknowledgments

We would like to thank the Goldrath lab and other co-authors for manuscript review. We thank the La Jolla Institute for Immunology Flow Cytometry Core facility and staff for their sorting assistance. We also thank the ImmGen consortium and ImmGenT team including David Zemmour and Ian Magill for their expertise and assistance with the pan-tissue scRNA-seq experiments. This work was performed in the context of the ImmGenT project within the ImmGen consortium, supported by R24072073. This work was supported by National Institutes of Health grants K00CA222711 (N.E.S.), GM133351 (A.R.), F31AI176705 (K.K.T), DP5OD031863 (B.K.), R35GM148339 (E.J.B.), R21AI178362 (A.W.G.), R01AI179952 (A.W.G.), R37AI067545 (A.W.G.), R01AI072117 (A.W.G.), and R01AI150282 (A.W.G.).

## Author contributions

Conceptualization: N.E.S and A.W.G. Methodology: N.E.S and A.W.G. Investigation: N.E.S., X.G., M.I.M., F.J., A.C., G.G., K.P.C., A.R., N.T., S.L.S., K.K.T., A.F., S.Q. Formal Analysis: N.E.S., X.G., M.I.M., M.H., A.M., G.G., S.L.S., M.A.B. Visualization: N.E.S., M.H., A.M., A.W.G. Data Curation: X.G., M.I.M., M.H., G.G., A.M., S.L.S. Resources: B.K., S.A.M., E.J.B., A.W.G. Writing – Original Draft: N.E.S. and A.W.G. Writing – Review & Editing: N.E.S., E.J.B., A.W.G. Supervision: B.K., S.A.M., E.J.B., A.W.G. Project Administration: B.K., S.A.M., E.J.B., A.W.G. Funding Acquisition: A.W.G.

---

**Figure S1:**
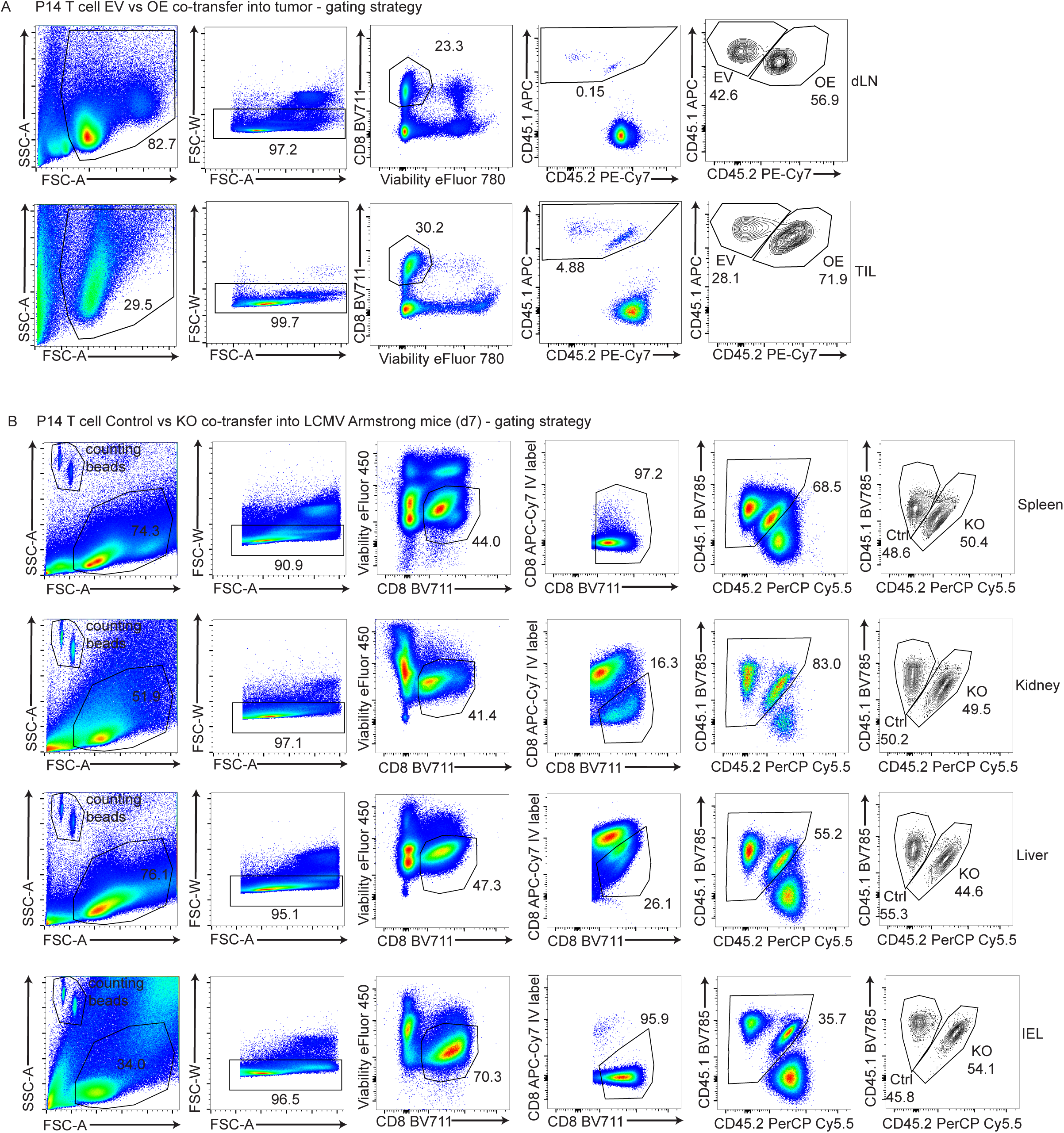
Gating Strategies for tumor and LCMV Armstrong. A) Gating strategy for EV vs ligase^OE^ P14 co-transfer experiments into tumor-bearing mice. B) Gating strategy for control vs ligase^KO^ P14 co-transfers into LCMV Armstrong-infected mice (d7 post-infection).

**Figure S2:**
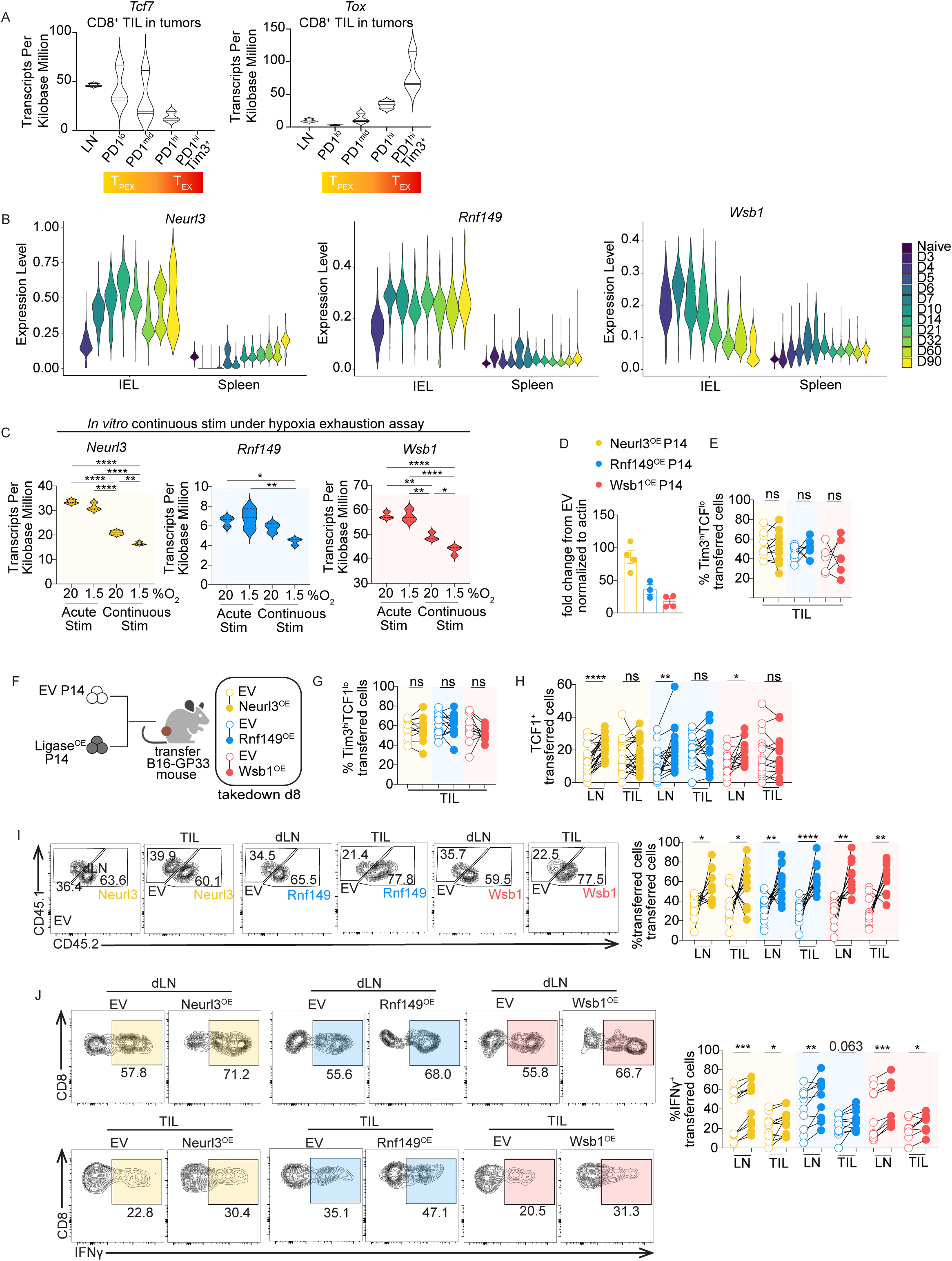
*Neurl3*, *Rnf149* and *Wsb1* expression in memory and exhaustion, and sustained ligase expression can improve TIL function in B16 melanoma, related to Figure 1. A) *Tcf7 or Tox* RNA TPM endogenous CD8^+^ T cells sorted from B16 melanoma dLN and TIL (GSE175408^33^). B) *Neurl3*, *Rnf149*, or *Wsb1* RNA expression in P14 T cells from IEL or spleens across infection timepoints (naive-d90) scRNA-seq (GSE131847^80^), quantified after MAGIC imputation. C) *Neurl3*, *Rnf149*, or *Wsb1* RNA TPM from CD8^+^ T cells under acute or continuous stimulation in hypoxic or normoxic environments (GSE155192^81^). D) qPCR validation of *Neurl3*, *Rnf149*, or *Wsb1* overexpression in P14 T cells (cDNA fold-change vs EV, normalized to actin) prior to adoptive transfer; four independent experiments. E) Tumor Tim3^hi^TCF1^lo^ P14 T cell frequencies of EV vs *Neurl3*^OE^, *Rnf149*^OE^, or *Wsb1*^OE^ 10 days after adoptive transfer into MC38-GP_33–41_ mice (*Neurl3*^OE^ n=10; *Rnf149*^OE^ n=8; *Wsb1*^OE^ n=6); two or more independent experiments. F) Experimental schematic: Congenically mismatched P14 T cells were activated and transduced 24 h later with Empty Vector (EV) or ligase-overexpressing retrovirus (Neurl3^OE^, Rnf149^OE^, or Wsb1^OE^). Next day, cells were sorted on ametrine reporter expression, cultured d5-6, then mixed 50:50 (1 × 10^6^ each, 2 × 10^6^ total) and transferred into d8 B16-GP_33–41_-bearing mice, dLN and TIL were analyzed 8 days later (BioRender). G) Transferred P14 T cell frequencies in dLN or tumors as in C (*Neurl3*^OE^ n=12; *Rnf149*^OE^ n=13; *Wsb1*^OE^ n=11); three independent experiments. H) Tumor Tim3^hi^TCF1^lo^ P14 T cell frequencies as in C (*Neurl3*^OE^ n=22; *Rnf149*^OE^ n=18; *Wsb1*^OE^ n=16); five independent experiments. I) TCF1^+^ P14 T cell frequencies as in C (*Neurl3*^OE^ n=22; *Rnf149*^OE^ n=18; *Wsb1*^OE^ n=16); five independent experiments. J) Representative flow plots and IFNɣ^+^ P14 T cell frequencies as in C followed by 4 h of peptide restimulation plus protein transport inhibitor (*Neurl3*^OE^ n=12; *Rnf149*^OE^ n=10; *Wsb1*^OE^ n=9); at least three independent experiments. Statistical significance was calculated using (C) one-way ANOVA, (D, G-J) paired Student’s *t*-tests. Connecting lines indicate that cells were isolated from the same mouse.

**Figure S3:**
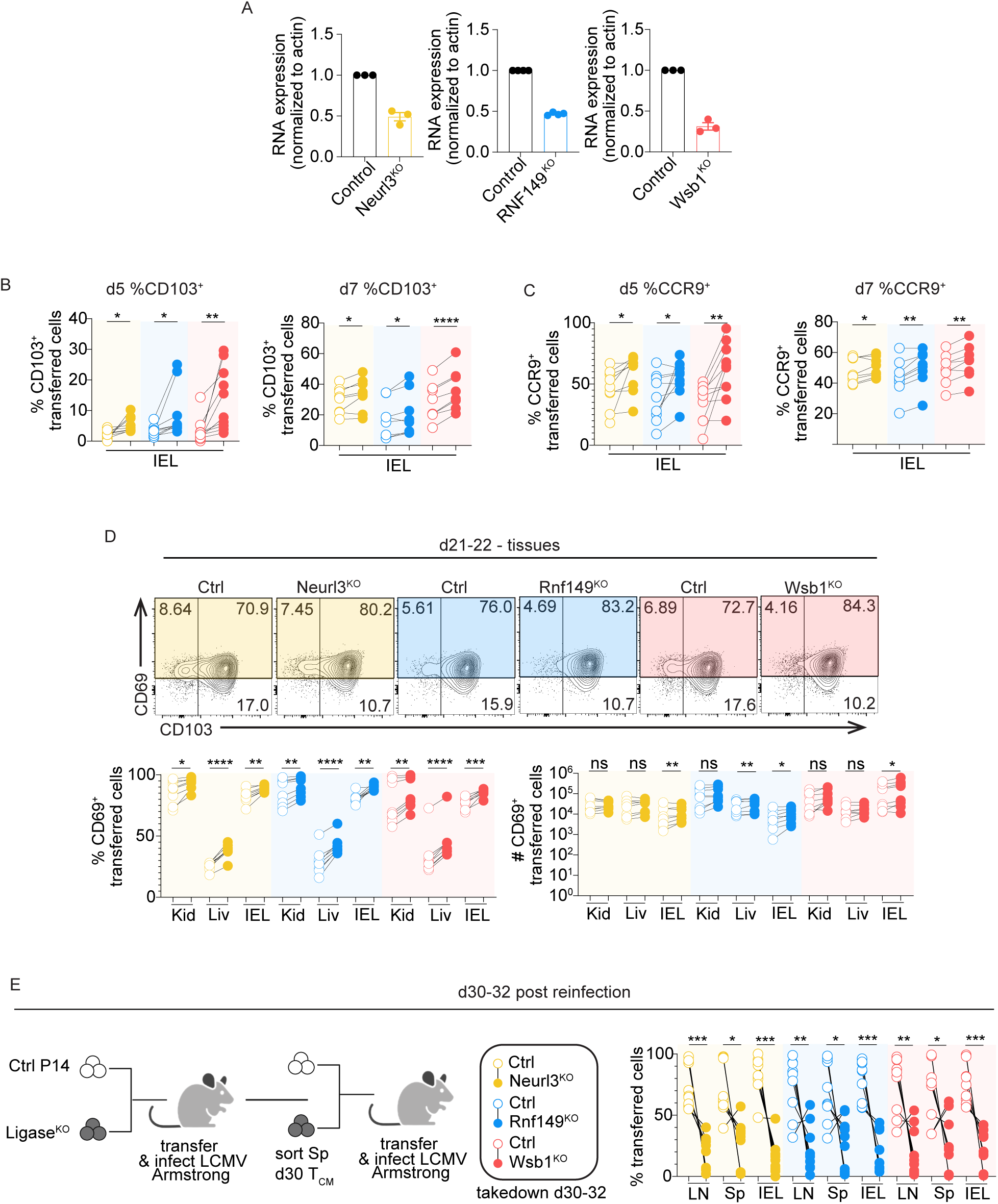
Loss of NEURL3, RNF149, or WSB1 expression accelerates d21-22 T_RM_ differentiation, related to Figure 3. A) qPCR validation of *Neurl3*, *Rnf149*, or *Wsb1* KO in P14 T cells (cDNA fold-change vs control, actin-normalized) prior to transfer; at least three independent experiments. B) CD103^+^ P14 T cell frequencies in the IEL of EV vs *Neurl3*^KO^, *Rnf149*^KO^, or *Wsb1*^KO^ 5 days (left) or 7 days (right) after transfer into LCMV Armstrong-infected mice (d5: *Neurl3*^KO^ n=9; *Rnf149*^KO^ n=9; *Wsb1*^KO^ n=10; for d7: *Neurl3*^KO^ n=8; *Rnf149*^KO^ n=8; *Wsb1*^KO^ n=8); two-three independent experiments. C) CCR9^+^ P14 T cell frequencies in the IEL of EV vs *Neurl3*^KO^, *Rnf149*^KO^, or *Wsb1*^KO^ 5 days (left) or 7 days (right) after transfer into LCMV Armstrong-infected mice (d5: *Neurl3*^KO^ n=9; *Rnf149*^KO^ n=9; *Wsb1*^KO^ n=10; d7: *Neurl3*^KO^ n=8; *Rnf149*^KO^ n=8; *Wsb1*^KO^ n=8); two-three independent experiments. D) CD69 vs CD103 in IEL flow plots and quantification of CD69^+^ P14 T frequencies and cell numbers in kidney, liver, and IEL with EV vs *Neurl3*^KO^, *Rnf149*^KO^, or *Wsb1*^KO^ 21–22 days post transfer into LCMV Armstrong-infected mice (*Neurl3*^KO^ n=8; *Rnf149*^KO^ n=8; *Wsb1*^KO^ n=8); two independent experiments. E) (Left) Serial-transfer schematic: Congenically mismatched P14 T cells were activated then nucleofected 24 h later with Thy1^KO^ Control CRISPR RNP, or ligase^KO^ (*Neurl3*^KO^, *Rnf149*^KO^, or *Wsb1*^KO^) plus Thy1^KO^ CRISPR RNP, mixed 50:50 (100,000 each; 200,000 total) and adoptively transferred into recipient mice that then received LCMV Armstrong. After 30 days, Ctrl and KO T_CM_ were sorted from LN and spleens, remixed 50:50 (6,000 total) and transferred into new recipients that then received LCMV Armstrong; tissues were analyzed d30-32, after IV labeling of circulating CD8^+^ T cells (BioRender). (Right) P14 T cell frequencies in the spleen, LN, and IEL of EV vs *Neurl3*^KO^, *Rnf149*^KO^, or *Wsb1*^KO^ 30-32 days after T_CM_ adoptive transfer into LCMV Armstrong-infected mice (*Neurl3*^KO^ n=9; *Rnf149*^KO^ n=9; *Wsb1*^KO^ n=9); three independent experiments. Statistical significance was calculated using (B-E) paired Student’s *t*-tests. Connecting lines indicate that cells were isolated from the same mouse.

**Figure S4:**
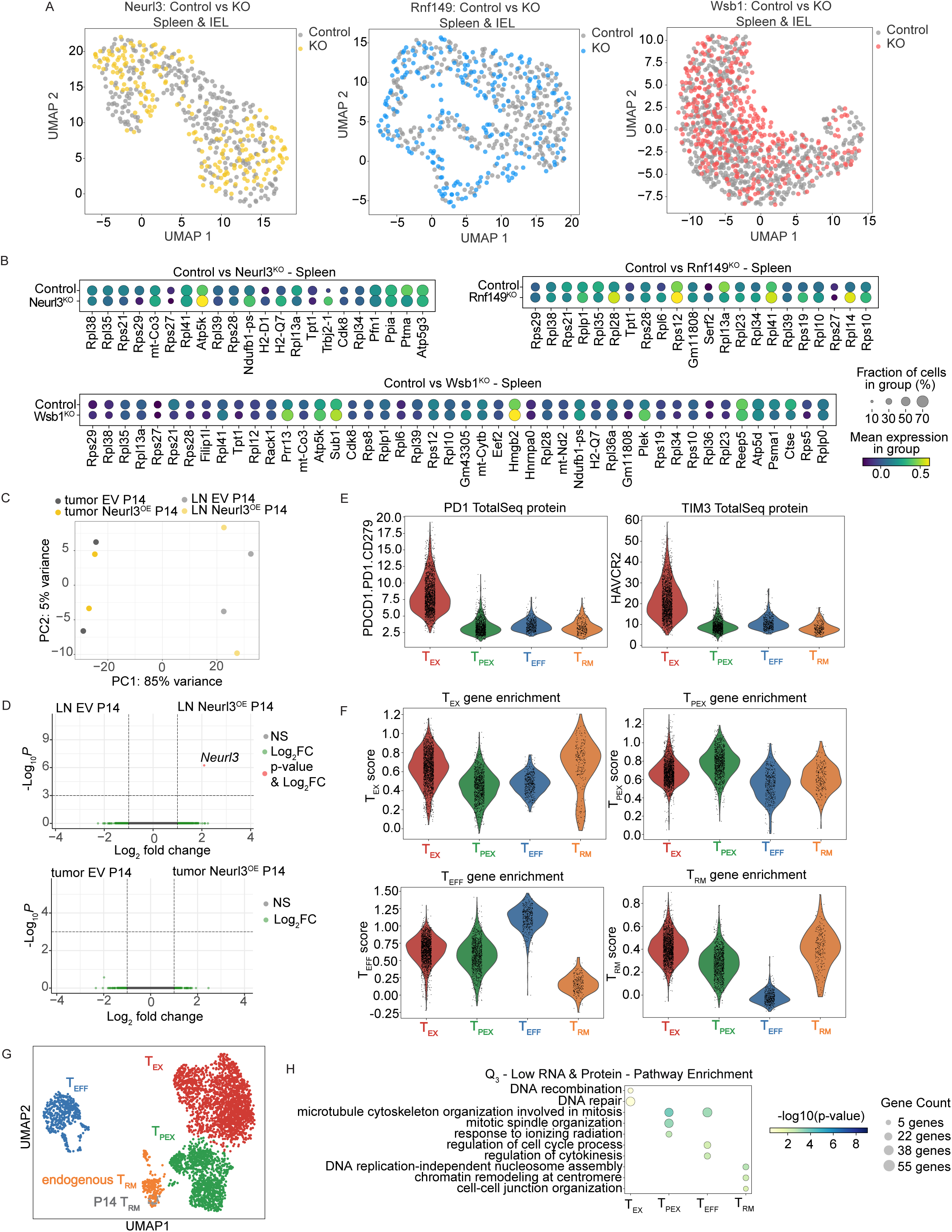
Transcriptomics of NEURL3, RNF149, or WSB1 KO or overexpression, and single cell RNAseq cluster characterization, related to Figure 3 and 4. A) RNA UMAP of control or ligase^KO^ P14 T cells isolated from the spleen or IEL 5 days after LCMV Armstrong-infection; four replicates per control-KO comparison in each tissue. B) Dot plot of splenic DE genes for control vs Neurl3^KO^ (top left), Rnf149^KO^ (top-right), and Wsb1^KO^ (bottom). Circle size = fraction of cells expressing gene, color = mean expression of gene. C) PCA of bulk RNA-seq from LN and tumor EV vs Neurl3^OE^ P14; n=2 pooled replicates per condition. D) Volcano plot from C: LN EV vs Neurl3^OE^ P14 (left) and tumor EV vs Neurl3^OE^ P14 (right). E) PD-1 (left) and Tim-3 (right) TotalSeq protein quantification across Figure 4 CITE-seq T cell groups. F) T_EX_, T_PEX_, and T_EFF_ T cell gene enrichment signatures^82^ and the T_RM_ gene enrichment signature^32^ overlaid on CITE-seq T cell clusters. G) UMAP as in Figure 4B, highlighting P14 IEL (gray) and endogenous T_RM_ (orange). H) Dot plot of genes from Q_3_ for each T cell subset (from Figure 4D), with GO:BP pathway enrichment for the top 3 significantly pathways per subset.

**Figure S5:**
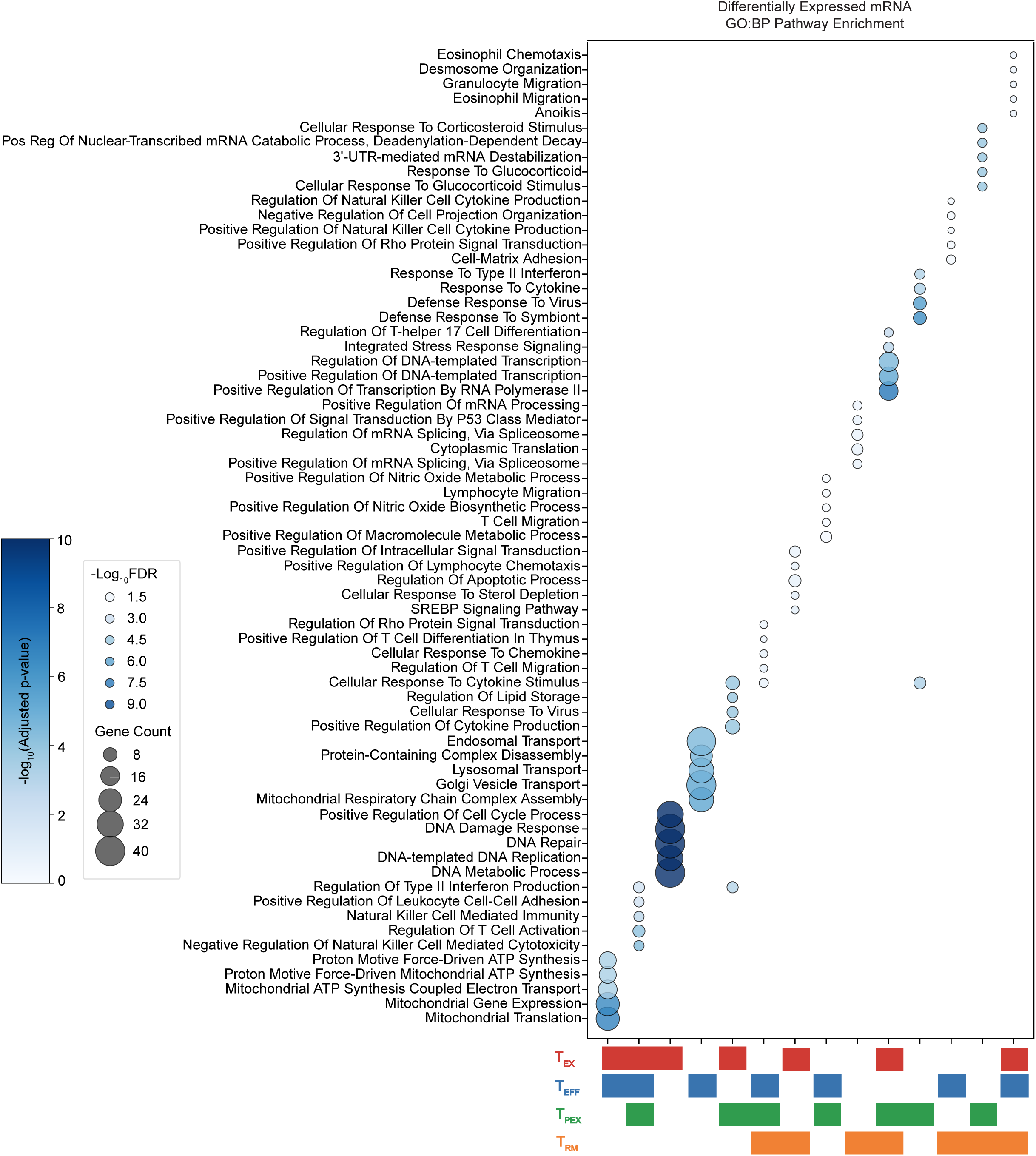
Single Cell RNAseq top unique and shared pathway enrichment, related to Figure 4. Bubble plot of GO:BP enrichment for differentially expressed RNA from Figure 4B.

**Figure S6:**
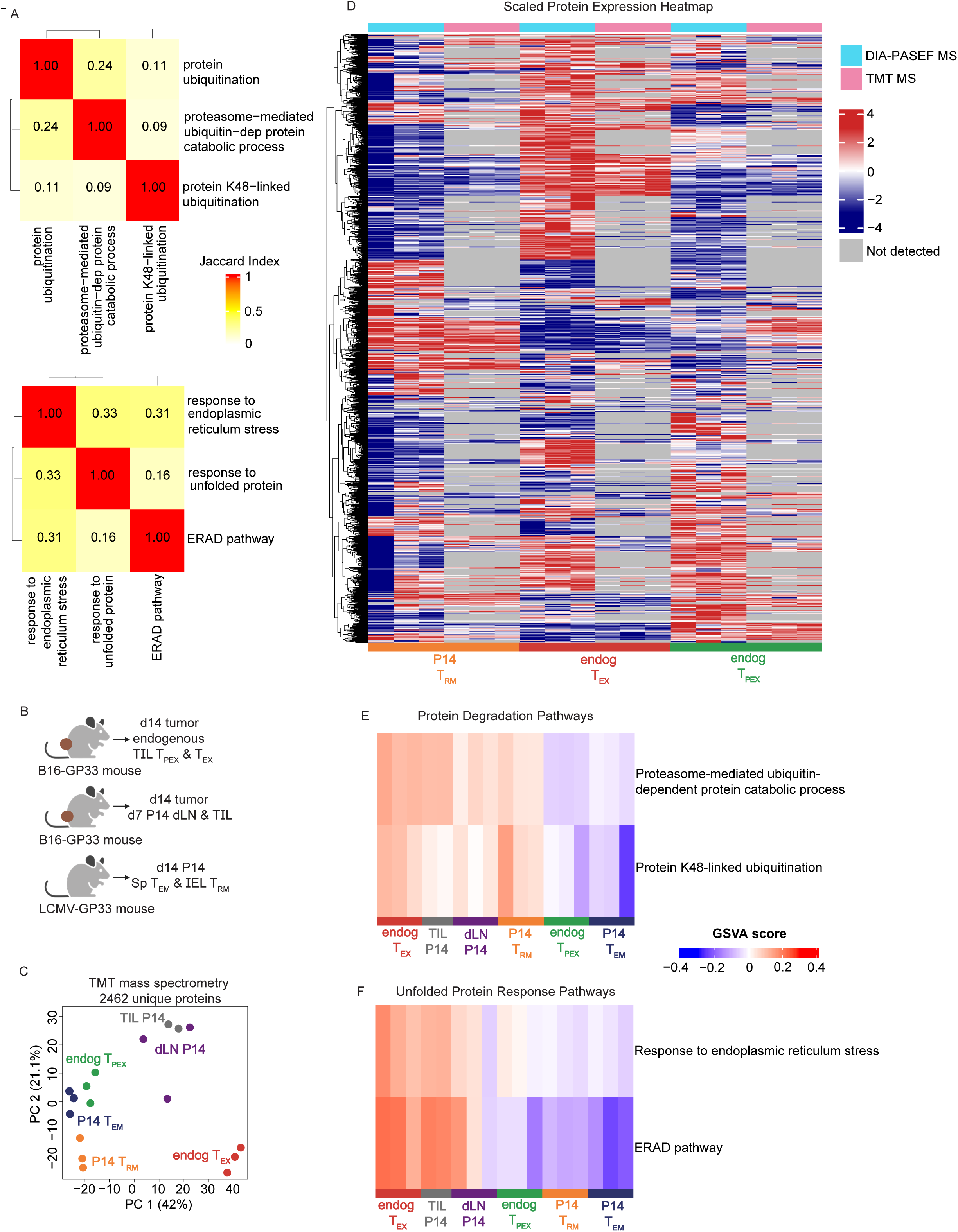
TMT mass spectrometry data of antigen-specific dLN and TIL, endogenous TIL, T_RM_ and T_EM_, related to Figures 4 and 5. A) Jaccard Index similarity among GO:BP protein-degradation (top) and unfolded protein response (bottom) terms; high Jaccard similarity (>0.7) indicates likely identical genes labeled with different GO terms while low values (0–0.3) indicates low gene overlap. B) Experimental schematic: CD8^+^ T cells were isolated from endogenous d14 B16 TIL (T_PEX_ sorted on CD8^+^PD1^lo/int^Tim3^−^ and T_EX_ sorted on CD8^+^PD1^hi^Tim3^+^), d14 B16 tumors that received *in vitro*-activated and expanded P14 (P14 transferred at d7), and from LCMV Armstrong memory P14 (d14 CD127^hi^CD62L^lo^ from the spleen (T_EM_) and P14 from the IEL (T_RM_)). (BioRender). C) TMT-MS PCA of CD8^+^ T cells from LCMV Armstrong memory P14 T_EM_ spleens and T_RM_, B16-GP_33–41_ P14 dLN and TIL from, and endogenous B16 T_PEX_ and T_EX_. Replicates: IEL n=3 (6 pooled/replicate), T_EM_ n=3 (6 pooled/replicate), P14 dLN n=3 (10 pooled/replicate), P14 TIL n=2 (10 pooled/replicate; one outlier removed from analysis), T_PEX_ n=3 (6 pooled/replicate), and T_EX_ n=3 (6 pooled/replicate). D) Scaled Protein Expression heatmap, of proteins shared between the DIA-PASEF mass spectrometry data (Figure 4C) and the TMT mass spectrometry data from B. Gray indicates the protein was not detected in the dataset. E) GSVA heatmap of protein degradation GO:BP pathways from C. F) GSVA heatmap of unfolded protein response GO:BP pathways from C.

**Figure S7:**
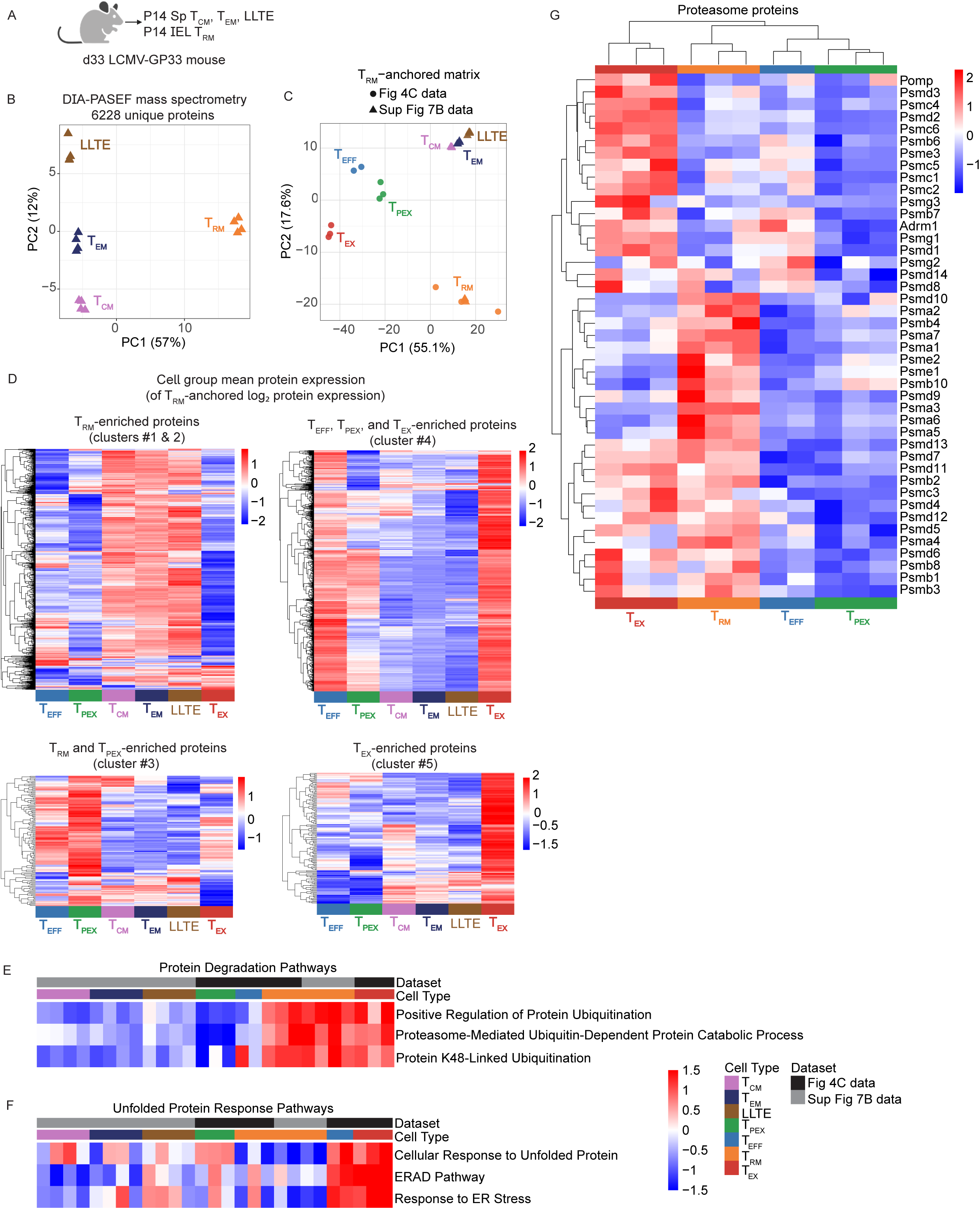
Circulating memory T cell proteomics, and proteasome of T_EFF_, T_RM_, T_PEX_, and T_EX_, related to Figures 4 and 5. A) Experimental schematic: naive P14 T cells were transferred and mice infected with LCMV Armstrong, obtained on d33 from the spleen (CD127^hi^CD62L^hi^ T_CM_, CD127^hi^CD62L^lo^ T_EM_, CD127^lo^CD62L^lo^ LLTE) and the IEL (T_RM_) (BioRender). B) Protein PCA of d33 from the spleen (CD127^hi^CD62L^hi^ T_CM_, CD127^hi^CD62L^lo^ T_EM_, CD127^lo^CD62L^lo^ LLTE) and the IEL (T_RM_), n=4 replicates (one mouse per replicate). C) PCA of the TRM-anchored, protein expression matrix combining Figure 4C data and Supplemental Figure 7B data. Protein values are log₂-transformed and T_RM_-centered within each dataset. D) Heatmaps of cell-group mean T_RM_-anchored log₂ protein expression for proteins in clusters 1+2, 3, 4, and 5 as defined in Figure 4G. T_RM_-anchored values were averaged within each group, and proteins with missing values were removed. E) GSVA heatmap of protein degradation GO:BP pathways, data from Figure Fig 4C and Supplemental 7B, combined using a T_RM_-anchored log₂ protein expression matrix. F) GSVA heatmap of unfolded protein response GO:BP pathways, data from Figure Fig 4C and Supplemental 7B, combined using a TRM-anchored log₂ protein expression matrix. G) Heatmap of proteasomal proteins from T_EFF_, T_RM_, T_PEX_, and T_EX_, expression normalized to row.

## Supplementary Tables

Supplementary Table 1: Core T_RM_ gene signature gene expression from endogenous CD8^+^ T cells sorted from B16 mouse melanoma dLN and TIL, related to figure 1.

> Bulk RNA-seq transcripts per kilobase million data derived from endogenous CD8+ T cells sorted from B16 mouse melanoma dLN and TIL (GSE175408), used to generate Figure 1A. RNAseq dataset was filtered on the core TRM gene signature.

Supplementary Table 2: Differentially expressed genes from d5 control and ligase^KO^ CITE-seq, related to figure 3.

> Pairwise RNA comparison of different T cell groups using the Wilcoxon rank-sum test.

Supplementary Table 3: Differentially expressed genes (pairwise comparisons) from *ex vivo* T_EFF_, T_RM_, T_PEX_, and T_EX_ CITE-seq, related to figure 4.

> Pairwise RNA comparison of different T cell groups using the Wilcoxon rank-sum test.

Supplementary Table 4: Differentially expressed proteins (pairwise comparisons) from *ex vivo* T_EFF_, T_RM_, T_PEX_, and T_EX_ DIA-PASEF mass spectrometry, related to figure 4.

> Pairwise protein comparison of different T cell groups using the Wilcoxon rank-sum test.

Supplementary Table 5: Differentially expressed protein cluster lists from *ex vivo* T_EFF_, T_RM_, T_PEX_, and T_EX_ DIA-PASEF mass spectrometry, related to figure 4.

> Lists of proteins per cluster, from Figure 4G.

Supplementary Table 6: Differentially expressed proteins (pairwise comparisons) from *ex vivo* TMT mass spectrometry, related to Supplemental Figure 6.

> Pairwise protein comparison of different T cell groups using the Wilcoxon rank-sum test.

Supplementary Table 7: Protein turnover list of short-lived proteins, related to figure 5.

> List of genes and corresponding half-lives (in hr).

## STAR methods

Reagents: See key resource table.

Software: See key resource table.

### Experimental Model and Study Participant Details

#### Mice

All mouse strains were bred and housed in specific pathogen-free conditions in accordance with the Institutional Animal Care and Use Guidelines of the University of California, San Diego (UCSD) in the temperature range of 18°–23°C in 40%–60% humidity. Male and female mice were used in the present study, between 6-12 weeks old. All the mice used had the C57BL/6J background. P14, Thy1.1 and CD45.1 congenic mice were bred in-house. All the animal studies were approved by the Institutional Animal Care and Use Committees of UCSD and performed in accordance with UC guidelines.

#### Cell culture

B16 melanoma cells and MC38 colorectal tumor cells that expressed the LCMV glycoprotein epitope amino acids 33–41 (B16-GP_33–41_) and PLAT-E cells were cultured in Dulbecco’s modified Eagle’s medium + D-glucose supplemented with 10% fetal bovine serum (FBS), 100 U ml^−1^ penicillin, 100 μg ml^−1^ streptomycin, 292 μg ml^−1^ L-glutamine, 1× non-essential amino acids, 1 mM sodium pyruvate, 10 mM HEPES and 55 µM 2-mercaptoethanol. The cell lines were confirmed to be free of mycoplasma using quantitative PCR (qPCR). Enriched CD8^+^ T cells were maintained in RPMI supplemented with 10% FBS, 100 U ml^−1^ penicillin, 100 µg ml^−1^ streptomycin, 292 µg ml^−1^ L-glutamine, 1× non-essential amino acids, 10 mM HEPES and 55 µM 2-mercaptoethanol. For the protein turnover experiment, T cells were cultured in SILAC media, including SILAC RPMI supplemented with 10% dialyzed FBS, 100 U ml^−1^ penicillin, 100 µg ml^−1^ streptomycin, 292 µg ml^−1^ L-glutamine, 1× non-essential amino acids, 10 mM HEPES and 55 µM 2-mercaptoethanol, heavy isotopes of arginine (^13^C_6_, ^15^N_4_) and lysine (^13^C_6_, ^15^N_2_), and 100 μM alanine^83^.

### Method Details

#### Tumor studies

For the MC38-GP_33–41_ TIL analysis, recipient mice were injected intradermally with 1 × 10^6^ MC38-GP_33–41_ adenocarcinoma cells. For co-transfers of transduced P14 cells, donor mice were sex-and age-matched to recipients or female donors, and the cells were transferred into recipients. On d5 post-activation, sorted ametrine^+^ cells were mixed at a 1:1 ratio and a total of 5 × 10^5^ P14 cells was transferred by intravenous (i.v.) injection into CD45 congenic recipients with d10 tumors. The mice were monitored three times per week, and the mice were sacrificed 10 days after the adoptive T cell transfer, for dLN and TIL analyses. For the MC38-GP_33–41_ tumor growth curves, a total of 1 × 10^6^ P14 cells was transferred by i.v. injection into CD45 congenic recipients with d8 tumors. The mice were monitored three times per week, and mice were removed from the study when their tumors extended 15 mm in any direction.

For the B16-GP_33–41_ TIL analysis, recipient mice were injected intradermally with 2.5 × 10^5^ B16-GP_33–41_ melanoma tumor cells. For co-transfers of transduced P14 cells, donor mice were sex-and age-matched to recipients or female donors, and the cells were transferred into recipients. On d5 post-activation, sorted ametrine^+^ cells were mixed at a 1:1 ratio and a total of 2 × 10^6^ P14 cells was transferred by i.v. injection into CD45 congenic recipients with d8 tumors. The mice were monitored three times per week, and mice were sacrificed 8 days after the adoptive T cell transfer, for dLN and TIL analyses. For the B16-GP_33–41_ tumor growth curves, a total of 5 × 10^6^ P14 cells was transferred by i.v. injection into CD45 congenic recipients with d6 tumors. Mice were also administered isotype or anti-PD-1 therapy, 200 μg of Armenian hamster isotype antibody or anti-PD-1 antibody intraperitoneally (i.p.) every other day. Mice were removed from the study when their tumors extended 15 mm in any direction.

#### Infection studies

Donor mice were sex- and age-matched to recipients or female donors, and the cells were transferred into recipients. For LCMV Amstrong experiments, C57BL/6J P14 CD8^+^ T cells congenic for CD45 or Thy1 were adoptively transferred at 2.5 × 10^4^ cells per recipient mouse by i.v. injection into CD45 or Thy1 congenic recipients. The mice were then infected with 2 × 10^5^ plaque-forming units (pfu) of LCMV Armstrong by i.p. injection 24 h after transfer. For co-transfers of nucleofected cells, P14 cells were mixed at a 1:1 ratio and a total of 5 × 10^5^ P14 cells was transferred by i.v. injection into CD45 or Thy1 congenic recipients. Thereafter, recipient mice were infected with 2 × 10^5^ pfu of LCMV by i.p. injection. For the experiment using sorted T_CM_ followed by reinfection, control or KO T_CM_ P14 cells mixed at a 1:1 ratio and a total of 6 × 10^3^ P14 cells was transferred by i.v. injection into CD45 or Thy1 congenic recipients. Thereafter, recipient mice were infected with 2 × 10^5^ pfu of LCMV by i.p. injection.

For LCMV Cl13 experiments, co-transfers of transduced P14 cells cells were mixed at a 1:1 ratio of sorted, ametrine^+^ cells, and a total of 2 × 10^3^ P14 cells was transferred by i.v. injection into CD45 or Thy1 congenic recipients. Thereafter, recipient mice were infected with 2 × 10^6^ pfu of LCMV by retro-orbital injection.

#### Preparations of single-cell suspensions (tumor)

Single-cell suspensions of lymphocytes were prepared by mechanical disaggregation followed by treatment with ACK (ammonium–chloride–potassium) lysing buffer. The tumor was minced into small pieces and then incubated in RPMI with 1.2 mg ml^−1^ of HEPES, 292 µg ml^−1^ L-glutamine, 1 mM MgCl_2_, 1 mM CaCl_2_, 5% FBS and 100 U ml^−1^ collagenase (Worthington) at 37°C for 30 min. Lymphocytes from the tumor were separated on a 44%/67% Percoll density gradient. For the B16 experiments, Percoll density gradient separation was followed by lymphocyte separation media to further purify the lymphocytes.

#### Preparation of single-cell suspensions (other non-lymphoid tissues)

To identify CD8^+^ T cells in the vasculature of nonlymphoid tissues (small intestine, kidney, and liver), 3 µg of anti-CD8α (clone 53-6.7) conjugated with APC-eFluor 780 were injected i.v. into the mice 5 min before sacrifice. Cells that displayed weak or no labeling with the anti-CD8α antibody were considered to be located outside the vasculature. Single-cell suspensions of splenocytes were prepared by mechanical disaggregation followed by treatment with ACK lysing buffer. Intraepithelial lymphocytes (IEL) were prepared through the removal of Peyer’s patches and the luminal contents. The small intestine was cut longitudinally and into 1-cm pieces, then incubated at 37°C for 30 min in Hanks’ balanced salt solution with 2.1 mg ml^−1^ sodium bicarbonate, 2.4 mg ml^−1^ HEPES, 8% bovine growth serum and 0.154 mg ml^−1^ DTE (EMD Millipore). The kidneys and liver were minced into small pieces and then incubated in RPMI with 1.2 mg ml^−1^ of HEPES, 292 µg ml^−1^ L-glutamine, 1 mM MgCl_2_, 1 mM CaCl_2_, 5% FBS and 100 U ml^−1^ collagenase (Worthington) at 37°C for 30 min. Lymphocytes from the small intestine, kidney, and liver were separated on a 44%/67% Percoll density gradient.

#### Generation of retrovirus-containing supernatants and CD8^+^ T cell transduction

To generate *Neurl3-*, *Rnf149-*, and *Wsb1*-overexpressing vectors, gBlocks of coding sequences (IDT) were cloned into MSCV-IRES-reporter vectors via NEBuilder HiFi DNA Assembly cloning, and then transformed into DH5alpha competent *Escherichia coli*. Successful cloning was confirmed by whole plasmid sequencing. Retroviral particles were produced by first plating 2 million PLAT-E cells on a 10-cm tissue culture dish 1 day before transfection. On the following day, each plate was transfected with 2 µg of pCL-Eco and 6 µg of the plasmid of interest using TransIT-LT1 (Mirus). After 24 h, plat-E media was replaced with T cell culture media to collect the retrovirus. On the same day, P14 T cells were isolated from the spleen and lymph nodes and negatively enriched; these cells were the resuspended with 2 µg/ml anti-CD28 antibody (catalog no. 37.51, eBioscience) and 50 U/ml IL-2 (CETUS) and plated in a 6-well dish that was coated with 5 µg/ml anti-CD3 antibody (catalog no. 145-2C11, eBioscience) for activation. Twenty-four hours after plating the T cells, retronectin-coated, non-tissue culture-treated plates were first incubated with retroviral media, after which the T cells resuspended in retroviral medium plus IL-2 were added to retronectin plates and spinfected for 40 min at 1,600 RPM at 37°C, followed by incubation overnight in retronectin plates with retroviral medium. Twenty-four hours after transduction, all the ametrine^+^ cells (as assessed by flow cytometry) were considered to be transduced. For the TIL analysis experiments, the cells were then sorted based on reporter expression, and expanded *in vitro* prior to adoptive transfer.

#### CRISPR-mediated gene knockout

P14 T cells were isolated from the spleen and lymph nodes and negatively enriched. They were then resuspended with 2 µg/ml anti-CD28 antibody (catalog no. 37.51, eBioscience) and 50 U/ml IL-2 (CETUS) and plated in a 6-well dish coated with 5 µg/ml anti-CD3 antibody (catalog no. 145-2C11, eBioscience) for activation. At 24 h post-activation, using the P3 Primary Cell 4D-Nucleofector X Kit and Unit, the T cells were electroporated with complexed tracrRNA (IDT), Cas9 (UC Berkeley), and pooled crRNA (IDT). crRNA targeting Thy1 (control) was used alone as a control and was mixed with the conditional crRNA to be used as an indicator of electroporated cells^84,85^.

#### Flow cytometry and cell sorting

For sorting, the cells were incubated with antibodies for 15 min at 4°C in PBS that was supplemented with 0.5% BSA and 2 mM EDTA, followed by cell collection in medium. For flow cytometry, the cells were incubated with the indicated antibodies for 15 min at 4°C in PBS that was supplemented with 2% bovine growth serum and 0.01% sodium azide. All cells were then fixed with Fixation Buffer (BioLegend) to preserve surface staining. Cytoplasmic staining was completed using the BD Cytofix/Cytoperm™ Fixation/Permeabilization Kit (BD), and intranuclear staining was performed using the FoxP3 transcription factor staining kit (eBioscience). For assays that involved CD8^+^ T cell stimulation, P14 cells were incubated for 4 h in T cell culture medium at 37°C with 10 nM GP_33–41_ peptide and Protein Transport Inhibitor (eBioscience), together with congenially distinct splenocytes for peptide presentation. K48 staining included 10 µM MG-132 and incubation at 37°C for 1 h prior to staining. Proteostat staining included congenically distinct, naive splenocytes added to each well for dye staining normalization, followed by surface staining, surface fixation, and cytoplasmic fixation, followed by staining with Proteostat dye (diluted 1:50,000 in perm wash). The dye was not washed out before running the samples, as according to manufacturer’s instructions. For ligase-overexpressing P14 Proteostat staining, Thy1.1 was used as a reporter, due to ametrine’s spectral overlap with Proteostat. For ImageStream visualization of Proteostat staining, the same numbers of cells per group were stained to normalize for background staining. Stained cells were analyzed using the LSRFortessa or LSRFortessa X-20 cytometers (BD), FACSDiva software, and Flowjo software (TreeStar). All sorting was performed on BD FACSAria Fusion instruments.

#### Real-Time PCR

RNA was isolated using the RNeasy Plus Mini Kit (QIAGEN). Complementary DNA was reverse-transcribed using the High Capacity cDNA Reverse Transcription Kit (Applied Biosystems). Real-Time PCR was performed with SYBR Green and primers for *Neurl3*, *Rnf149*, *Wsb1*, and *Actb*; quantification was performed using the *ΔΔC*^t^ method.

#### Proteasome activity assay

Single-cell suspensions of LN and TIL were sorted as described above. After sorting, 5,000 cells per well were plated in 100 µl of medium in a white-walled, clear-bottom plate, with or without 10 µM MG-132 and incubated at 37°C for 1 h. Proteasome activity was then assessed using the Cell-Based Proteasome-Glo 3-Substrate System (Promega) according to the manufacturer’s instructions.

#### CITE-seq of CD3ε^+^ T cells from B16-GP_33–41_ and LCMV Armstrong

##### Experimental setup, staining, and sorting – B16-GP_33–41_

Recipient mice were injected intradermally with 5 × 10^5^ B16-GP_33–41_ melanoma tumor cells. Congenically distinct CD45.1.2 P14 T cells and CD45.1 Pmel T cells were activated in vitro (plate-bound CD3 and CD28), expanded to day 5, then 2.5 × 10^6^ of each transgenic were transferred into day 7 tumor recipients (CD45.2 B6 mice). Mice also began 200ug anti-4-1BB + 50ug anti-PD-1 checkpoint blockade immunotherapy or isotype control on the day of adoptive T cell transfer, then every other day thereafter. CD3ε+ T cells from dLN or TIL were profiled by CITE-seq on day 7 after transfer. We first sorted (unless otherwise specified) 90,000 endogenous CD3ε+ T cells or 11,250 transferred T cells from the dLN or tumor into DMEM supplemented with 2% bovine growth serum and 10 mM HEPES. Due to cell number limitations, 10,000 endogenous CD3ε+ cells, 6000 P14 T cells, and 1000 Pmel T cells were sorted from the immunotherapeutic tumor group. When enough cells could not be sorted from the original sample, unstained cells were adding to get a total 500,000 cells. Following the first sort, hash-tagged cells were pooled incubated with the ImmGenT TotalSeq-C custom mouse panel containing a custom panel of 128 antibody-derived tags (ADT; see STAR methods) and washed thoroughly. For the second sort, 50,000 CD3ε+ T total cells from the pooled tissues were sorted. 5,000 CD3ε+ T cells from a naive spleen were also sorted to allow for batch correction with totalVI.

##### Experimental setup, staining, and sorting – LCMV-Armstrong

Prior to LCMV infection, congenically distinct 5 x 10^4^ naive CD45.1 P14 T cells and 1 x 10^5^ naive CD45.1.2 SMARTA T cells were transferred into naive CD45.2 B6 recipients. Then, CD3ε+ T cells from siIEL. siLPL, prostate, SG, lung, bone marrow, spleen, mesenteric lymph nodes and mediastinal lymph nodes were profiled by CITE-seq on days 7, 30, and 60 of LCMV Armstrong infection. For each timepoint, we first sorted 5x10^4^ CD3ε+ T cells from the spleen, prostate, SG, bone marrow, mesenteric lymph node, siIEL, siLPL, lung and mediastinal lymph node were initially sorted (unless otherwise specified) into DMEM supplemented with 2% bovine growth serum and 10 mM HEPES. Due to cell number limitations, on day 7 4.36 x10^3^ CD3e cells were sorted from the SG and 4.32 x10^3^ CD3e cells from the prostate. On day 30, 25,106 CD3e cells from the SG, 19,277 CD3e cells from the prostate and 1,245 CD3e cells were sorted from the lung. When 5x10^4^ cells could not be sorted from the original sample, unstained cells were adding to get a total 5x10^4^ cells. Following the first sort, hash-tagged cells were pooled incubated with ImmGenT TotalSeq-C custom mouse panel containing a custom panel of 128 antibody-derived tags (ADT; see STAR methods) and washed thoroughly. For the second sort, 5x10^4^ CD3ε+ T total cells from the pooled tissues were sorted. For each timepoint, 5x10^3^ CD3ε+ T cells from a naïve spleen were also sorted to allow for batch correction with totalVI.

##### Single-cell RNA, TCR, and TotalSeq-C sequencing—Library preparation

The collected cells were pooled and split into two technical replicates (independent libraries) for all subsequent steps. scRNA-seq was performed using 10X Genomics 5’v2 with feature barcoding for cell surface protein and immune receptor mapping following the manufacturer’s instructions. After cDNA amplification, the smaller fragments containing the TotalSeq-C–derived cDNA were purified and saved for feature barcode library construction. The larger fragments containing transcript-derived cDNA were saved for TCR and gene expression library construction. For both cDNA portions, the library size was measured with an Agilent Bioanalyzer 2100 high-sensitivity DNA assay and quantified using a Qubit dsDNA HS assay kit on a Qubit 4.0 fluorometer.

TCR cDNA was amplified from the transcript-derived cDNA by nested PCR according to the manufacturer’s protocol. The TCR libraries were then subjected to enzymatic fragmentation, ligation and indexing with unique Dual Index TT set A index sequences with an index PCR. Transcript-derived cDNA was processed into RNA libraries. After enzymatic fragmentation and size selection, purified cDNA was ligated to an Illumina R2 sequence and indexed with unique Dual Index TT set A index sequences with an index PCR. TotalSeq-C–derived cDNA was processed into feature barcode libraries. The cDNA was indexed with unique Dual Index TN set A index sequences with an index PCR. The three libraries were pooled together on the basis of molarity with relative proportions of 47.5% RNA, 47.5% feature barcode, and 5% TCR. This pool was then sequenced on an Illumina NovaSeq S2, 100c. The sequencing configuration followed 10X Genomics specifications.

##### Single-cell RNA, TCR, and TotalSeq-C sequencing—Data processing

Gene and TotalSeq-C antibody (surface protein panel and hashtags) counts were obtained by aligning reads to mm10(GRCm38) and the M25(GRCm38.p6) GeneCode annotation (GRCm38.p6) of the mouse genome and the DNA barcode for the TotalSeq-C panel using CellRanger software (v7.1.0) with default parameters. Cells were distinguished from droplets with high RNA and TotalSeq-C counts using inflection points on a total count curve (barcodeRanks function in the DropletUtils package). Samples were demultiplexed using hashtag oligo (HTO) counts and the HashSolo algorithm^86^, and doublets (cells positive for multiple hashtags) were excluded. Surface protein expression was normalized using the DSB method to remove ambient background and technical noise^87^. Cells were filtered if they met any of the following criteria: expression of two isotype control antibodies, fewer than 150 RNA UMI counts, or >5% mitochondrial gene expression. RNA counts were normalized to 10,000 transcripts per cell using normalize_total() and log-transformed with log1p in Scanpy^88^. Highly variable genes were selected using the Seurat v3 method with batch correction^89^, and expression data were scaled to unit variance (max value = 10) prior to integration. To jointly model RNA and protein data, we applied totalVI from the scvi-tools framework^90^. RNA and protein modalities from 3 conditions (d7 LCMV, d30 LCMV, Tumor) were concatenated using MuData, and totalVI was trained to a shared latent space accounting for batch effects and noise.

##### Clustering, dimensionality reduction, and subset annotation

Cells of interest were first subset from the larger multimodal dataset based on protein expression of CD4 and CD8A. TotalVI-denoised protein values for CD4 and CD8A were visualized in a 2D scatter plot, and thresholds were manually defined to retain CD8-enriched and CD4-low cells, isolating CD8⁺ T cell populations. To further refine the dataset, P14 TCR transgenic T cells were identified by high co-expression of TCRVA2 and CD45.1 proteins, using denoised TotalVI protein layers. For RNA-based clustering, we used the totalVI-derived RNA embedding (X_pca) and computed a neighborhood graph using 10 nearest neighbors (sc.pp.neighbors). Clustering was performed using the Leiden algorithm^91^ at a resolution of 0.1. UMAP was used for visualization (sc.tl.umap). Clusters were annotated post hoc based on enrichment of previously defined transcriptional signatures for T_RM_^10^, T_PEX_, T_EX_, and T_EFF_^3^ subsets. Enrichment scores were calculated using scanpy.tl.score_genes() for each signature and violin plots were generated to visualize signature distribution across clusters. TIL T_PEX_ vs T_EX_ clusters were also confirmed using the protein modality PDCD1 and HAVCR2 expression.

##### Differential expression and pathway analysis

For each unique pairwise comparison between clusters, a Wilcoxon rank-sum test was applied using scanpy.tl.rank_genes_groups() to identify significantly differentially expressed genes (DEGs)^88^ by RNA expression. Genes were considered significant if they had an adjusted p-value < 0.05 and an absolute log₂ fold-change > 1.0. DEGs across clusters was used to construct gene “signatures,” representing unique combinations of clusters in which a gene was upregulated. Each signature was stored as a tuple and linked to its associated genes, enabling identification of genes uniquely or jointly expressed across subsets. Each gene signature was subjected to enrichment analysis using the mouse GO:Biological Process 2023 gene set via Enrichr in Python through the gseapy package, with background set to genes expressed in the dataset^92^. Signatures with fewer than five genes were excluded. For each valid group the top five pathways (by adjusted p-value) were tagged, all significantly enriched pathways (FDR < 0.05) were retained, and combined pathway lists were exported, both globally and per T cell subset (T_RM_, T_PEX_, T_EX_, T_EFF_). To visualize pathways enriched across gene signatures, a bubble plot was generated using Seaborn and Matplotlib^93,94^. The size of each point reflected the number of genes contributing to enrichment, while color intensity encoded statistical significance as –log₁₀(FDR), capped at 10. Pathway names were plotted on the y-axis and gene signature combinations on the x-axis. GSVA was performed on normalized scRNA-seq data using defined clusters. Differentially expressed genes between clusters were first identified with Wilcoxon test, and the union of significant DEGs (|log₂FC| > 1, FDR < 0.05) was used. GO Biological Process gene sets were retrieved with gseapy (GO_Biological_Process_2023, Mouse), mapped to mouse gene symbols, and GSVA scores were computed per cell; pathway differences across clusters were tested by one-way ANOVA with Benjamini–Hochberg FDR correction, and significant pathways were visualized as GSVA score heatmaps using seaborn and matplotlib.

#### CITE-seq of d5 ligase^KO^ P14 T cells from LCMV Armstrong

##### Experimental setup, staining, and sorting

Congenically distinct P14 T cells were activated and electroporated with complexed tracrRNA, Cas9, and crRNA (as described above) to generate CRISPR control and KO P14 T cells, which were co-transferred into recipients then infected with LCMV Armstrong. On day 5 of infection, P14 T cells from IEL, liver, and spleen, were profiled by CITE-seq. We first sorted 5x10^4^ P14 T cells from the IEL, liver, and spleen of 4 pooled mice per group (control vs NEURL3^KO^, control vs RNF149^KO^, and control vs WSB1^KO^) into DMEM supplemented with 2% bovine growth serum and 10 mM HEPES. Due to the number of groups, all control samples were pooled together and all KO samples were pooled together. Following the first sort, hash-tagged cells were pooled incubated with antibody-derived tags (ADT, TotalSeq™-C Mouse Universal Cocktail, V1.0) and washed thoroughly. For the second sort, all stained cells from the pooled tissues were sorted as well as 5x10^4^ CD8+ T cells from a naive spleen were also sorted to allow for batch correction with totalVI.

##### Single-cell RNA and TotalSeq-C sequencing—Library preparation

scRNA-seq was performed using 10X Genomics 5’v2 with feature barcoding for cell surface protein and immune receptor mapping following the manufacturer’s instructions. After cDNA amplification, the smaller fragments containing the TotalSeq-C–derived cDNA were purified and saved for feature barcode library construction. The larger fragments containing transcript-derived cDNA were saved for gene expression library construction. For both cDNA portions, the library size was measured with an Agilent Bioanalyzer D5000 high-sensitivity DNA assay and quantified using a Qubit 2.0 fluorometer.

Transcript-derived cDNA was processed into RNA libraries. After enzymatic fragmentation and size selection, purified cDNA was ligated to an Illumina R2 sequence and indexed with unique Dual Index TT set A index sequences with an index PCR. TotalSeq-C–derived cDNA was processed into feature barcode libraries. The cDNA was indexed with unique Dual Index TN set A index sequences with an index PCR. The two libraries were pooled together on the basis of molarity with relative proportions of 40% GEX (RNA) and 10% surface (feature barcode) for each control and KO pooled samples. This pool was then sequenced on an Illumina NovaSeq X Plus 10B. The sequencing configuration followed 10X Genomics specifications.

##### Single-cell RNA and TotalSeq-C sequencing—Data processing

Gene expression and TotalSeq-C antibody (surface protein panel and hashtags) counts were obtained using CellRanger (v7.1.0, 10x Genomics) with default parameters. Reads were aligned to the *mm10* (2020) reference transcriptome and the DNA barcode for the TotalSeq-C panel. Samples were demultiplexed using hashtag oligo (HTO) counts and the HashSolo algorithm^86^, and doublets (cells positive for multiple hashtags) were excluded. Surface protein expression was normalized using the DSB method to remove ambient background and technical noise^87^. Cells were filtered if they met any of the following criteria: expression of two isotype control antibodies, fewer than 150 RNA UMI counts, or >5% mitochondrial gene expression. RNA counts were normalized to 10,000 transcripts per cell using normalize_total() and log-transformed with log1p in Scanpy^88^. Highly variable genes were selected using the Seurat v3 method with batch correction^89^, and expression data were scaled to unit variance (max value = 10) prior to integration. To jointly model RNA and protein data, we applied totalVI from the scvi-tools framework^90^. RNA and protein modalities from 2 conditions (Control and KO) were concatenated using MuData, and totalVI was trained to a shared latent space accounting for batch effects and noise.

##### Dimensionality reduction & differential expression

Dimensionality reduction was performed using contrastiveVI^95^. UMAP projections were computed using the salient representation, min_dist=1.5. Differentially expressed (DE) genes between knockout (KO) and control samples were identified using the Wilcoxon rank-sum test as implemented in Scanpy’s rank_genes_groups() function^88^, DE genes were considered significant if they had an adjusted p-value < 0.05 and an absolute log fold change > 0.1, and visualized using dot plots.

#### EV and *Neurl3^OE^* P14 – dLN and TIL bulk RNAseq

Recipient mice were injected intradermally with 1 × 10^6^ MC38-GP_33–41_ adenocarcinoma cells. Congenically distinct P14 T cells were activated and transduced (as described above) to generate EV and *Neurl3^OE^* P14 T cells, which were co-transferred into recipients on d5 post-activation cells were mixed at a 1:1 ratio and a total of 5 × 10^5^ P14 cells was transferred by intravenous (i.v.) injection into CD45 congenic recipients with d6 tumors. The mice were monitored three times per week, and the mice were sacrificed 6 days after the adoptive T cell transfer. For each replicate, cells from 2-3 mice were pooled per tissue and then 5,000 P14 sorted for bulk RNA-seq. 1 × 10^3^ were then resorted into 1× TCL lysis buffer + 1% 2-mercaptoethanol. Library preparation for ultra-low-input RNA-seq was performed as described online (https://www.immgen.org/img/Protocols/ImmGenULI_RNAseq_methods.pdf).

Trimmomatic was used to remove adapters and trim low-quality reads (NexteraPE-PE.fa:2:30:10:1:TRUE LEADING:3 TRAILING:3 SLIDINGWINDOW:4:15 MINLEN:25)^96^. Trimmed reads were then aligned to the gencode M25 annotation of the mm10 genome using STAR with the default conditions^97^. Aligned reads were then quantified with featureCounts^98^ (-t exon -g gene_id -p -B) and DEG was identified using DEseq2^99^. PCA analysis in was generated with the plotPCA function in DEseq2. Volcano Plot was generated with EnhancedVolcano.

#### Analysis of human melanoma single-cell RNA-seq dataset

Single-cell RNA-seq data from human melanoma patients treated with checkpoint blockade^60^ was processed beginning with raw cell-by-gene matrices and associated metadata. Cells were filtered to retain those expressing at least 200 genes, and genes were retained if they were detected in three or more cells. Normalization was performed using Scanpy’s normalize_total and log1p functions^88^. Highly variable genes were identified and used for PCA dimensionality reduction. A k-nearest neighbors graph was constructed in PCA space, followed by UMAP embedding^100^ and Leiden clustering^91^. Clusters expressing CD8 markers were identified and retained for downstream analysis, and only cells with CD8A expression above a defined threshold were kept. These CD8⁺ cells were re-clustered, inspected, and clusters of interest were identified based on expression profiles of canonical markers. A subset of clusters were extracted and relabeled (e.g., T_PEX_, T_EX_, T_RM_-like-TIL, etc.) based on gene expression in clusters. UMAP was recomputed for this subset and used for visualization. These renamed clusters were treated as categorical variables and used to stratify subsequent expression analyses. A heatmap was generated using canonical markers and the E3 ligases *RNF149* and *WSB1* to illustrate overlap with transcriptional T cell subsets, and expression values were scaled between 0 and 1 for comparability across genes. Excluding cells with zero expression, violin plots were then generated to visualize *RNF149* and *WSB1* expression across CD8⁺ T cells, stratified by treatment and response status, and significance was tested using unpaired t-test.

#### Analysis of T_RM_ infection time course and T_RM_ across tissues

For analysis of the P14 infection time course^80^ and P14 T_RM_ across tissues^78^, each were combined into a counts matrix using the merge function in Seurat. Data were log(normalized) and scaled using NormalizeData and ScaleData. Data imputation was performed using MAGIC^101^ with the log(normalized expression) values and the default settings and the exact solver and plotted as violin plots.

#### Data Independent Acquisition-parallel accumulation–serial fragmentation mass spectrometry

For the T_EFF_, T_PEX_, T_EX_, and T_RM_ dataset, naive P14 were adoptively transferred into mice and infected with LCMV as described above. At d7, 2 replicates of P14 from the spleen (T_EFF_) and at d14, 3 replicates comprised of 6 pooled samples of P14 from the IEL (T_RM_) were isolated, washed with PBS, and frozen as cell pellets. D14 was used as an early memory timepoint due to cell number limitations at d30. For endogenous TIL, mice received B16-GP_33–41_ tumors as described above. Ad d14, 3 replicates comprised of 6 pooled samples each for T_PEX_, and 3 replicates comprised of 6 pooled samples each for T_EX_ (T_PEX_ sorted on CD8^+^PD1^lo/int^Tim3^−^ and T_EX_ sorted on CD8^+^PD1^hi^Tim3^+^) were isolated, washed with PBS, and frozen as cell pellets.

For the T_CM_, T_EM_, LLTE, and T_RM_ dataset naive P14 were adoptively transferred into mice and infected with LCMV as described above. At d33, 4 replicates of P14 from the spleen (CD127^hi^CD62L^hi^ T_CM_, CD127^hi^CD62L^lo^ T_EM_, CD127^lo^CD62L^lo^ LLTE) and the IEL (T_RM_) were isolated. The sample processing approach for low input T cell proteomics was adapted from a previous study^102^. 2,000 T cells were sorted into a PCR tube containing 2 μL of freshly prepared cell lysis buffer (0.045% n-dodecyl-β-D-maltoside, 90 mM TEAB pH 8.5) and snap frozen on dry ice.

##### Sample preparation (T_EFF_, T_PEX_, T_EX_, and T_RM_)

Cell pellets, each containing 10^5^ sorted cells, were resuspended in 50 μL of freshly prepared urea lysis buffer (8 M urea, 75 mM NaCl, 50 mM Tris-HCl, pH 8.0, 1 mM NaF, 1 mM sodium orthovanadate, 1 mM β-glycerophosphate). The suspension was sonicated for 300 s (30 s on/30 s off cycles) in a water bath sonicator (Fosmon, model #51028HOM). Protein was reduced with 10 mM TCEP for 30 min, alkylated with 15 mM freshly prepared iodoacetamide (IAA) for 45 min in the dark, and then treated with 10 mM DTT for 15 min to quench excess IAA. The samples were diluted to a final volume of 400 μL with 50 mM Tris-HCl, pH 8.0. For digestion, 200 ng Lys-C and 400 ng sequencing grade modified trypsin were added to the samples, which were then incubated at 37 °C with shaking overnight. The digestion was terminated by adding formic acid to achieve a final concentration of 1%.

##### Sample preparation (T_CM_, T_EM_, LLTE, and T_RM_)

The samples were incubated at 95°C for 5 min and then at 4°C for 1 min in a thermal cycler (Applied Biosystems). For digestion, 1 μL of 100 ng sequencing grade modified trypsin was added to the solution, which was then incubated at 37 °C in a thermal cycler overnight. The digestion was terminated by adding 1 μL of 6% formic acid.

##### Analysis (T_EFF_, T_PEX_, T_EX_, and T_RM_)

The digests were desalted using the Stage-Tip method. Subsequently, 2 μL of the 10 μL peptide sample was analyzed using an LC-MS/MS setup comprising a timsTOF Pro 2 mass spectrometer (Bruker Daltonics) coupled to a nanoElute 2 nano-LC system (Bruker Daltonics) via a CaptiveSpray ion source. Peptides were first loaded onto a trap column (PepMap Neo, 5 mm x 300 μm Trap) and then eluted through a reversed-phase C18 column (PepSep, 25 cm x 150 μm, 1.5 μm) maintained at 50 °C over a 50 min gradient of 5-35% solvent B (acetonitrile in 0.1% formic acid) at a flow rate of 450 nL/min. The peptides were analyzed using data-independent acquisition (dia) Parallel Accumulation-Serial Fragmentation (PASEF) mode. The isolation windows for dia-PASEF were determined based on data-dependent acquisition (dda)-PASEF data^103^ and generated by implementing the py_diAID algorithm to maximize proteome coverage^104^. The settings for dda-PASEF included the following: a ramp time and an accumulation time of 75 ms each, one MS scan, and 10 PASEF MS/MS ramps per acquisition cycle. The MS survey scan covered 100-1700 m/z and 0.6-1.6 V⋅s/cm². Precursors with up to 5 charges were selected, with an active exclusion time of 0.4 min. For dia-PASEF, 50 variable isolation windows for 25 dia-PASEF scan cycles (two windows per cycle) were designed to encompass the precursor distribution across the m/z-1/K0 plane as defined by the dda-PASEF data, ranging from 323.6 to 1221.6 m/z and 0.7 to 1.34 V⋅s/cm², resulting in a cycle time of 2.11 s. Collision energy was decreased linearly with increasing mobility, starting from 59 eV at 1/K0 = 1.6 V⋅s/cm² and reducing to 20 eV at 1/K0 = 0.6 V⋅s/cm².

##### Analysis (T_CM_, T_EM_, LLTE, and T_RM_)

The peptides were directly loaded onto a trap column (PepMap Neo, 5 mm x 300 μm Trap), desalted online with 20 μL of solvent A (0.1% formic acid in H2O), and then eluted through a reversed-phase C18 column (PepSep, 25 cm x 150 μm, 1.5 μm) maintained at 50 °C over a 50 min gradient of 2-35% solvent B (0.1% formic acid in acetonitrile) at a flow rate of 300 nL/min. The dia-PASEF was performed as described above with the following modifications: high-sensitivity detection was enabled; ion accumulation and ramp time were set to 200 ms; each MS1 full scan was followed by 10 dia-PASEF MS2 scans (two isolation windows per scan), resulting in a cycle time of 2.2 s.

##### Data processing (T_EFF_, T_PEX_, T_EX_, and T_RM_)

The dia-PASEF raw files were processed using a library-free search approach in DIA-NN version 1.8.1^105^. An in silico digestion of a UniProt Mus musculus reference proteome (downloaded in January 2023) was employed to predict the spectral library. The search parameters applied included the following: tryptic digestion with up to two missed cleavages, carbamidomethylation of cysteine as a static modification, and oxidation of methionine and N-terminal acetylation as variable modifications. The precursor false discovery rate (FDR) threshold was set at 1%. For quantification, the quantification strategy was set to Robust LC (high precision) and cross-run normalization was set to RT-dependent. Match-between-runs (MBR) was enabled. The resulting protein group matrix from DIA-NN, reported at a 1% FDR (global q-values for protein groups; global and run-specific q-values for precursors), was used for downstream and statistical analysis. The statistical analysis was performed using the limma package in R^106^.

##### Data processing (T_CM_, T_EM_, LLTE, and T_RM_)

The data processing was performed as described above with the following modifications: DIA-NN version 2.2 was used; cysteine modification was unchecked; the quantification strategy was set to QuantUMS (high precision).

##### Data analysis – differentially expressed proteins and pathway enrichment

Principal component analysis (PCA) was performed on variance-stabilized data (vst and rlog transformations) to visualize sample clustering. Data were structured as a count matrix for differential expression analysis using the DESeq2 package^99^. Sample metadata were loaded in parallel and matched to sample columns for experimental group assignment. Differential expression analysis was performed using a negative binomial generalized linear model (GLM) framework implemented in DESeq2, with group.type used as the design formula. For each pairwise group comparison, significantly differentially expressed proteins were defined as those with an adjusted *p*-value < 0.05 and an absolute log₂ fold change > 0.5.

Heatmaps of normalized expression values for the union of all DEGs were generated using the pheatmap package. To explore functional patterns among DEGs, hierarchical clustering was applied to the DEG genes were partitioned into 9 expression clusters. Gene Ontology (GO) enrichment analysis for Biological Process (BP) terms was performed on each cluster using the clusterProfiler package^107^, with gene symbol-to-Entrez ID mapping via org.Mm.eg.db. Top enriched mouse GO terms were visualized using a custom bubble plot showing –log₁₀ (adjusted *p*-value) and number of genes per term, with background set to proteins expressed in the dataset. Redundant GO terms were collapsed by prioritizing terms associated with shared gene sets and lower adjusted *p*-values. All visualizations were created using ggplot2.

GSVA was used to calculate sample-wise enrichment scores using log-transformed, variance-stabilized intensity values. Mouse GO:BP gene sets were used as input, and enrichment scores were computed using the Gaussian kernel density estimation method (kcdf=“Gaussian”) in the GSVA package^108^. Differential enrichment of GSVA scores across groups was assessed using the limma package^109^, and significantly enriched pathways (FDR < 0.05) were identified via linear modeling and empirical Bayes moderation. Pathways related to proteasomal degradation and the unfolded protein response (UPR) were selected based on biological relevance and filtered to remove redundant GO terms, and heatmaps were generated via clustered cell type. Redundancy and overlap among enriched gene sets were assessed via Jaccard similarity matrices.

##### T_RM_-anchored normalization and integration

Proteomic data from the two DIA-PASEF datasets were processed in R: protein intensities were log₂-transformed, and sample-median centered. Within each dataset, TRM samples were used as an internal reference to compute a per-gene TRM mean, which was subtracted to generate a TRM-centered matrix. TRM-centered matrices from both datasets were then combined on the union of genes, and genes with any non-finite values were removed for downstream analyses.

##### PCA, heatmaps, GSVA and pathway analysis

PCA was performed in R using prcomp (base R) and visualized with ggplot2. For heatmaps, TRM-anchored log₂ values were summarized as cell group mean protein expression on a TRM-anchored log₂ scale, row-z-scored, and plotted with pheatmap, using DE and LFC of 0.5-filtered gene clusters (1–5) as defined in Figure 4G. GSVA was performed with the GSVA R package on the TRM-anchored matrix using mouse GO:BP gene sets to generate pathway activity scores across T cell groups to identify significantly enriched protein degradation and unfolded protein response pathways.

#### RNA vs protein analysis - RNA–Protein graph, quadrant-based pathway enrichment analysis, and distance matrix

To assess the concordance between gene expression at the transcriptomic and proteomic levels, pseudobulk RNA-seq data and averaged protein intensities were merged by gene symbol. Scatter plots comparing RNA and protein abundances were generated, and genes were assigned to one of four quadrants based on their log-transformed expression relative to the geometric mean of RNA and protein values. For each quadrant (Q1–Q4), gene ontology enrichment analysis was performed using Enrichr via the gseapy package, and the top 3 biological process (GO:BP) pathways (filtered on ≥5 gene expression) were extracted per quadrant, with background set to genes expressed in the dataset ^92^. The enrichment results were visualized using quadrant-wise bubble plots. Dot sizes encoded the number of genes per pathway and color indicated statistical significance (–log₁₀ *p*-value).

To compare global transcriptome- and proteome-level similarity across subsets, principal component analysis (PCA) was conducted separately for RNA and protein data using scanpy^88^. Pairwise Euclidean distances between subsets in PCA space were computed using scipy.spatial.distance_matrix, and results were visualized as heatmaps.

#### Tandem Mass Tag mass spectrometry

For memory T cells, naive P14 were adoptively transferred into mice and infected with LCMV as described above. At d14, 3 replicates comprised of 6 pooled samples of CD127^hi^CD62L^lo^ P14 from the spleen (T_EM_) and P14 from the IEL (T_RM_) were isolated, washed with PBS, and frozen as cell pellets. D14 was used as an early memory timepoint due to cell number limitations at d30. For endogenous TIL, mice received B16-GP_33–41_ tumors as described above. Ad d14, 3 replicates comprised of 6 pooled samples each for T_PEX_, and 3 replicates comprised of 6 pooled samples each for T_EX_ (T_PEX_ sorted on CD8^+^PD1^lo/int^Tim3^−^ and T_EX_ sorted on CD8^+^PD1^hi^Tim3^+^) were isolated, washed with PBS, and frozen as cell pellets. For P14 dLN and TIL, mice received B16-GP_33–41_ tumors as described above, then received d5 *in vitro*-activated P14 on d7 after tumor implantation. On d14, 3 replicates comprised of 10 pooled samples each for P14 dLN, 2 replicates comprised of 10 pooled samples each for P14 TIL were isolated, washed with PBS, and frozen as cell pellets.

TMT-based proteomics was performed as previously described^45^. Briefly, samples with several modifications. First, TMTpro reagents were lysed, reduced, and alkylated using 8 M urea, 10 mM TCEP, and 10 mM iodoacetamide. Urea was then diluted to <2 M using 50 mM ammonium bicarbonate, and sequencing-grade trypsin was added at a 1:100 enzyme-to-substrate ratio utilized for overnight digestion at 37°C. The entire digest was passed through a Stage-tip and the on-tip labeling was performed using TMTpro reagents by adding 1 µL of reagent to 100 µL of freshly prepared. 1 uL of TMTpro was added to 50 mM phosphate buffer (pH 8.0), which was then passed before passing over the C18-bound peptides. Peptides were fractionated using Stage-tip-based SDB-RPS fractionation. Peptides were eluted stepwise using increasing concentrations of acetonitrile (5–90%) in 20 mM ammonium formate, pH 10. Fractions were vacuum-dried and stored at –80°C until LC-MS analysis. Mass spectrometry data were tip. Data was acquired usingon an Orbitrap Eclipse (Thermo Scientific) mass spectrometer.

Raw data were processed using Spectrum Mill software (Agilent and Broad Institute) and searched against the UniProt Mouse database. Carbamidomethylation of cysteine and TMTpro on lysines and peptide N-termini were set as fixed modifications, with oxidation of methionine and protein N-terminal acetylation set as variable modifications. Up to three missed tryptic cleavages were allowed. Peptide and protein FDRs were controlled at <1% using a target-decoy approach. Protein quantification was performed using TMT reporter ion intensities normalized to a pooled reference channel. Proteins were reported if identified by at least two unique peptides and a protein score ≥20. Only one representative proteoform per gene was used for downstream analyses).

GSVA was used to calculate sample-wise enrichment scores using log-transformed, variance-stabilized intensity values. Mouse GO:BP gene sets were used as input, and enrichment scores were computed using the Gaussian kernel density estimation method (kcdf=“Gaussian”) in the GSVA package^108^. Differential enrichment of GSVA scores across groups was assessed using the limma package^109^, and significantly enriched pathways (FDR < 0.05) were identified via linear modeling and empirical Bayes moderation. Pathways related to proteasomal degradation and the unfolded protein response (UPR) were selected based on biological relevance and filtered to remove redundant GO terms, and heatmap was generated via clustered cell type.

To compare T cell proteomic signatures across the two mass spectrometry datasets, intensity values were log₂-transformed (where applicable) and rescaled per gene to the range [–3, 3] using min–max normalization to account for differences in dynamic range across studies. Combined heatmaps were constructed by merging the scaled matrices from both studies. Rows were clustered using hierarchical clustering on row-wise imputed matrices (mean imputation for missing values), while columns were manually ordered to reflect biological groupings. Heatmaps were generated using the pheatmap package and included a top annotation bar to indicate dataset origin. Missing values were colored in gray to distinguish from scaled intensities.

#### Protein turnover mass spectrometry

##### Experimental design

WT T cells were activated and electroporated with complexed tracrRNA, Cas9, and crRNA (as described above) to generate CRISPR control and KO T cells, which were then switched to T_RM_ culture conditions (25 U/mL IL-2, 10 ng/mL IL-15, 10 ng/mL TGFβ and 10 nM retinoic acid)^110^ 1 day after nucleofection. Cells are expanded 2 more days in T_RM_ culture conditions, then on day 5 post-activation, cells were counted, washed with PBS, and frozen as pellets (for timepoint 0 hr), then remaining cells were switched to SILAC media including heavy isotopes of arginine (^13^C_6_, ^15^N_4_) and lysine (^13^C_6_, ^15^N_2_) plus T_RM_ cytokines. Cells were then sampled, counted, washed with PBS, and frozen as pellets for 2, 4, 8, 12, 16, 24, 32, 40, 48, 72 hr.

##### Sample Preparation

Cell pellets were lysed in protein (RIPA) lysis buffer (10mM Tris-Cl pH 8.0; 1.5mM EDTA; 1% Triton X-100; 0.1% SDS; 140 mM NaCl; 0.1% sodium deoxycholate; phosphatase and protease inhibitor cocktails (ThermoFisher Scientific)). Lysates were incubated on ice for 30min, sonicated for 10 seconds at 40% amplitude, and centrifuged at 14000 rcf for 10 minutes at 4°C. The supernatant was collected, and protein concentration was determined using Pierce BCA Protein Assay Kit (ThermoFisher Scientific) and a Molecular Devices SpectraMax iD3. A total of 30µg of protein was reduced with TCEP (ThermoFisher Scientific) to a concentration of 10mM and alkylated with IAA (Sigma) to a concentration of 30mM. Peptides underwent chloroform/methanol extraction (Fisher Chemical) and digested overnight at 47°C with sequencing grade modified porcine trypsin (Promega) in Triethylammonium bicarbonate (TEAB, Honeywell Fluka) at a protein:trypsin ratio of 50:1. The following day, samples were acidified to 0.5% formic acid/0.1% trifluoroacetic acid (TFA, Fisher Chemical). After plate equilibration with Buffer B, tryptic peptides were loaded onto a Sep-Pak C18 96-well plate (Waters) to be desalted using a Resolvex A200 (Tecan) automated method where samples underwent four washes with Buffer A. Peptides were eluted from the plate using 40% Buffer B. After equilibrating the plate with 70% Buffer B, samples were filtered using an AcroPrep Advance Filter Plate (Pall). Samples were lyophilized overnight (SpeedVac SPD210, ThermoFisher Scientific) and stored at -80°C. Buffer A = 0.1% formic acid, 0.5% acetonitrile in water, Buffer B = 80% acetonitrile, 20% water, 0.1% formic acid.

##### Analysis

SILAC LC-MS analysis was conducted using an EvoSep One (EvoSep) coupled to a trapped ion mobility spectrometry quadrupole time-of-flight mass spectrometer (timsTOF Pro, Bruker Daltonik) by a captive spray ion source (Bruker Daltonik). Tryptic peptides were reconstituted in Buffer A. 500 ng of sample was loaded on C18 Evotips (Evosep) according to the manufacturer’s protocol. Peptides were separated on a 15cm x 150μm ReproSil-pur C18 Endurance analytical column with 1.9 µm beads (EvoSep) at 50°C using a linear 44 min gradient 30 SPD method (Evosep). Mass spectrometry analysis was performed in data-independent (dia-PASEF) mode with 1 MS1 survey TIMS-MS and 10 PASEF MS/MS scans per acquisition cycle. The captive spray ion source capillary voltage was set to 1650 V. Both the ramp time and ion accumulation time were set to 100.0 ms. The ion mobility ranged from 1/K0 = 0.85 to 1.27 V⋅s/cm², excluding singly or triply charged precursor ions. Full scan mass ranged from 100 to 1700 m/z. Collision energy was decreased linearly with increasing mobility, starting from 59 eV at 1/K0 = 1.6 V⋅s/cm² and reducing to 20 eV at 1/K0 = 0.6 V⋅s/cm².

##### Data Processing

Raw files acquired from abundance LC-MS analysis were analyzed with Spectronaut software (version 18.2) using the directDIA method with MS2 scans used for protein and peptide quantification. The UniProt murine proteome reference (Mouse_UP000000589_10090) containing 21,756 protein entries was used with protein and peptide-level false discovery rates (FDR) set to 0.01. Selected variable modifications included N-terminal acetylation and methionine oxidation while cysteine carbamidomethylation was set as a fixed modification. Trypsin was selected as the digestion enzyme with a maximum of two allowed missed cleavages and a minimum peptide length of seven amino acids. A total of 90,392 peptides and 7,520 protein groups were identified. Protein MS2 exclusive intensity values were quality-assessed using ProteiNorm^111^. Data normalization was performed with Variance Stabilizing Normalization (VSN)^112^, followed by statistical analysis using proteoDA^113^. Linear Models for Microarray Data (limma) combined with empirical Bayes (eBayes) smoothing were applied to the standard errors^109^. Proteins with a false discovery rate (FDR)-adjusted p-value of less than 0.05 were considered statistically significant.

Raw files acquired from SILAC LC-MS analysis were analyzed with Spectronaut software (version 20.3). Using the Pulsar search engine, a DIA-based spectral library was created using the “Generate Library from Pulsar/Search Archives” function. The UniProt murine proteome reference (Mouse_UP000000589_10090) containing 21,756 protein entries was used with protein and peptide-level false discovery rates (FDR) set to 0.01. Selected variable modifications included N-terminal acetylation and methionine oxidation while cysteine carbamidomethylation was set as a fixed modification. Trypsin was selected as the digestion enzyme with a maximum of two allowed missed cleavages and a minimum peptide length of seven amino acids. To ensure robust filtering in multiplexed SILAC DIA data, the Multi Channel Q-value Filter was set to “Channel Qvalue,” enforcing channel-specific confidence estimation. A total of 92,824 peptides and 7,604 protein groups were identified. A labeled spectral library was then generated, utilizing two label channels with Arg10 and Lys8 modifications added to the second channel. Raw files were searched using the labeled spectral library with MS2 scans used for protein and peptide quantification and half-life determination.

##### Protein half-life determination

The statistical software R (v4.4.3) was used to calculate protein half-life. Peptide-level quantification were processed to compute the percentage of light-labeled peptides and organized by protein, gene, condition, replicate. Peptides were filtered to include only those that contained at least 11 timepoints between 0 and 72 hours and excluding those with an increasing light fraction over time (suggesting incomplete labeling or biological noise). Using the fitdecay1 function from the R package nlfitr (v0.1.0) a non-linear leas squares (NLS) decay model was fit for each peptide time course, with upper and lower asymptotes fixed at 1 and 0 respectively, and weighted fitting enabled. The R package broom (v1.0.7) was used to extract the model output. Peptides with valid model fits were evaluated for goodness-of-fit by calculating pseudo and adjusted R-squared values. Only peptides with an adjusted R-squared ≥ 0.9 were retained. These peptides were averaged across proteins to compute mean percent light curves that were re-fit using the same NLS procedure to obtain protein-label kinetic parameters. The decay constant (k) was converted to half-life for each protein and condition (half-life = log2(k)).

##### Short-Lived Protein Abundance Analysis

To compare the abundance of short-lived (half-life ≤8 h) versus non-short-lived (half-life >8 h to ≤20 h) proteins across T cell populations, protein annotations for short-lived proteins were curated into a list using half-life quantification from either of the two replicates. Then protein intensity values obtained from the DIA-PASEF dataset were used to evaluate the abundance of short-lived proteins across T cell populations. Proteins were annotated as “short-lived” or “non-short-lived” based on curated list. Intensities were summed for each protein within a sample. For each biological replicate, the total abundance was computed as the sum of intensities for all detected proteins within each category. Mean abundance and standard error were calculated across replicates using the dplyr package. Visualization of group-wise means and significance was performed with ggplot2 and ggpubr, using bar plots with error bars and annotated *p*-values. For rank–abundance analysis, each T cell population was converted from protein intensities to within-sample proportions, averaged these across replicates, and ranked proteins by mean proportional contribution to plot faceted rank–abundance curves (implemented in R with dplyr, tidyr, and ggplot2). For stacked composition plots, the top 20 proteins per population were visualized (and the remaining collapsed as “Other”) as 100% stacked bars with ordered fills (implemented with dplyr, forcats, ggplot2, and scales). For Protein Set Enrichment Analysis, peptide-count matrices values were log2-transformed (log2[count+1]) in R. Pairwise differential ranking statistics between group.type levels were computed using limma (linear modeling with empirical Bayes moderation), and a short-lived proteins set was tested for enrichment against the ranked t-statistics using fgsea (fgseaMultilevel). GSEA-style running enrichment score plots were generated with ggplot2.

#### Quantitative and Statistical Methods

Statistical tests were performed using Prism (7.0/9.0; GraphPad), R (v.4.1), and Python (3.11.9) in VS Code software packages. Two-tailed, paired or unpaired Student’s *t*-tests, or one- or two-way analyses of variance (ANOVA) were used for comparisons between groups. *P*-values <0.05 were considered statistically significant. Values of P < 0.05 were ranked as follows: *, P < 0.05; **, P < 0.01; ***, P < 0.001; and ****, P < 0.0001.

